# CD4⁺ T cells confer transplantable rejuvenation via Rivers of telomeres

**DOI:** 10.1101/2025.11.14.688504

**Authors:** Alessio Lanna, Salvatore Valvo, Michael L. Dustin, Federica Rinaldi

**Affiliations:** Sentcell UK laboratories, Tuscany Life Sciences, GSK Vaccine Campus, Siena. Italy; Kennedy Institute of Rheumatology, Nuffield Department of Orthopaedics, Rheumatology and Musculoskeletal Sciences, University of Oxford, Oxford, UK; University College London, Division of Medicine, London. United Kingdom

## Abstract

The role of the immune system in regulating organismal lifespan remains poorly understood. Here, we show that CD4⁺ T cells release “telomere Rivers” into circulation after acquiring telomeres from antigen-presenting cells (APCs). River formation requires fatty acid oxidation at the T cell–APC synapse, which selectively excludes glyceraldehyde 3-phosphate dehydrogenase (GAPDH) from the telomere vesicles. The resulting Rivers are depleted of glycolytic enzymes but enriched in T cell–derived stemness factors, enabling targeted rejuvenation of senescent tissues across multiple organs. In aged mice, adoptive transfer of young or metabolically reprogrammed CD4⁺ T cells triggered River production *in vivo*, and Rivers isolated from these animals could be transplanted into other aged mice to propagate the rejuvenation phenotype independently of T cells. River therapy extended median lifespan by ∼17 months, with several mice surviving to nearly five years. This immune-driven telomere transfer pathway is conserved across kingdoms, including plants, defining the first systemic, transplantable programme of youth.

## Main

The search for a “Fountain of Youth” began long ago. Ageing is associated with chronic functional decline, disease, and mortality. Although its causes remain incompletely understood, modern studies suggest that blood — or blood-derived factors — can exert tissue-specific healing and rejuvenating effects, as seen in parabiosis^1,2^. However, these early findings have been challenged^3^, and no definitive mechanism for systemic rejuvenation has been established. One theory attributes ageing to the accumulation of terminally differentiated or senescent cells in multiple tissues, disrupting homeostasis ^4^. A true fountain of youth would need to target senescent cells across organs, be tightly regulated, and transfer youth-promoting activity from a young organism to an old one — as in the original parabiosis studies.

One rejuvenation candidate arises from telomere transfer between immune cells ^5,6^. We previously showed that antigen-presenting cells (APCs) donate telomere-containing vesicles to CD4⁺ T cells during immune synapse formation, extending their telomeres, preventing senescence, and generating long-lived, stem-like memory T cells ^5,6^. However, although fatty acid oxidation (FAO) supports the survival of memory T cells ^7,8^, it is not known whether FAO is required for telomere transfer — and whether this process could initiate a rejuvenation programme beyond the immune system itself. Here we show that, after telomere acquisition, recipient CD4⁺ T cells undergoing FAO, assemble and release “Rivers” of telomeres into the circulation. These Rivers recycle surplus APC telomeres unused by the T cells and rejuvenate tissues throughout the body, extending lifespan — an unprecedented programme in which CD4⁺ T cells transmit youth-promoting signals between organisms.

While analysing antigen-specific T cell memory responses, we observed that APC telomere transfer was accompanied by abundant extracellular telomeric material decorating CPT1A^+^ memory T cells ^7,8^ in OVA-reactive lymph nodes (CPT1A shuttles long-chain fatty acids into mitochondria during FAO sustaining long-lived memory T cells; **Fig. 1a**), but not in naïve mice (**Fig. 1a bottom**; Supplementary information 1). Histology revealed that these extracellular telomeres were not merely tethered to T cells but arranged in vessel-like networks (**Fig. 1b**), suggesting release into circulation. The elongated, punctate structures appeared to flow along these networks, evoking miniature streams of genetic material — henceforth referred to as telomere Rivers.

**Figure 1:**
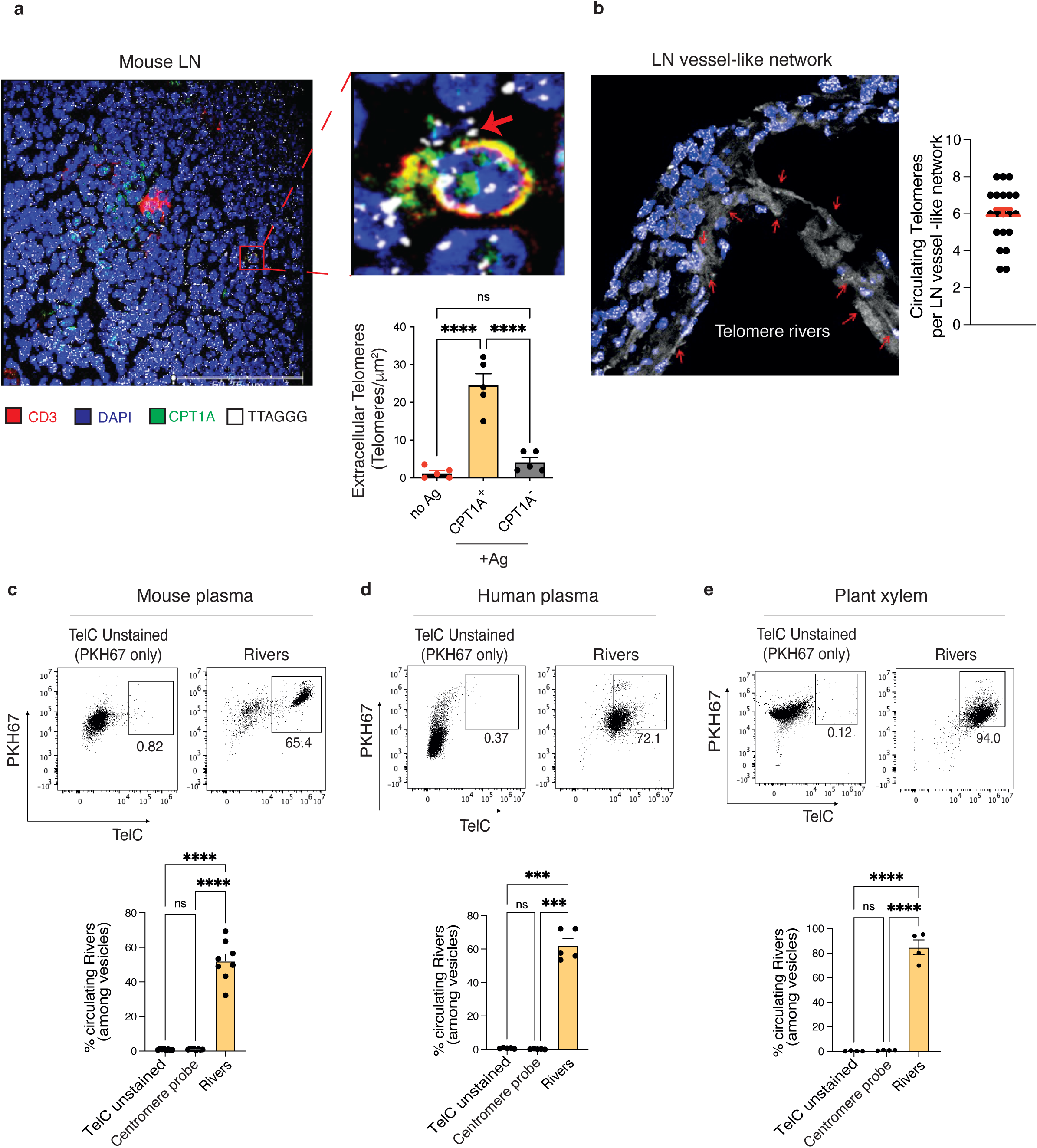
Discovery of telomere Rivers. (**a, left**) Extracellular telomeres *in vivo*. Reactive lymph-nodes were harvested from 6-week-old mice, 7 days after recall responses to ovalbumin then analyzed by IF-FISH staining (telomere probe) plus CD3 and Cpt1a antibodies. Nuclei staining demonstrated by DAPI. Representative of 3 separate mice. Scale bar, 50 μM. (**a, middle**) Magnification of extracellular telomeres as in (a; scale bar, 2 μM) and quantification of extracellular telomeres in Cpt1a^Bright^ and Cpt1a^Low^ (DMFICpt1a> 23%) T cells. Note lack of extracellular telomeres in T cell with lower Cpt1a expression. (**a, right**) Presence of extracellular telomeres in lymph node vessels, with examples denoted by red arrows, throughout. Scale bar, 50 μM. Three unchallenged animals were used as control (no Ag, Supplementary information 1). (**b**) Discovery of telomere release in efferent vessels of lymph nodes of mice vaccinated twice with ovalbumin (OVA; 30 µg) then analyzed 7 days after the recall response. Representative of 3 separate mice. DAPI, blue; TelC, white. Quantification of River per LN section is shown. Representative FACS profiles and quantifications of circulating telomere rivers in mouse (**c**; *n* = 8 mice) and human (**d**, *n* = 5 donors) sera and in plant xilema (**e**, *n* = 4 orchids). PKH67, lipid dye and TelC probe were used for telomere River identification across kingdoms. For quantification, each dot is an individual donor. Note that centromere probes had no detection (see also Supplementary information 1). One-way Anova with Bonferroni post-correction for multiple comparisons in (**a**) and unpaired Student’s t-test (**c-e**). ****P < 0.0001. Error bars indicate s.e.m.

In plasma from immunised mice, flow cytometry with TelC probes and PKH67 membrane dyes revealed abundant circulating telomeric vesicles (**Fig. 1c**), absent in negative controls and centromere-DNA probes. Vesicle TZAP retention^6^ confirmed telomere identity (Supplementary information 2). Similar structures were detectable in human plasma (**Fig. 1d**) and in plant xylem sap (**Fig. 1e**), where River formation required photosynthesis-driven β-oxidation ^9^ and was restored by the CPT1A activator fenofibrate ^10,11^ in darkness (**Fig. 1e**; Supplementary Information 3). Thus, extracellular telomere Rivers occur across kingdoms and appear following antigen-driven T cell responses.

Plant data suggested that FAO might also be required for River formation in immune cells. We therefore examined human CD27⁻CD28⁻CD4⁺ senescent T cells (T_sen_) ^12,13^, which lack CPT1A (ED Fig. 1a), divert fatty acids into ceramide accumulation ^14^, and — critically — fail to acquire telomeres from APCs^6^. This defect arises from impaired TCR clustering ^15^ at the immune synapse (ED Fig. 1b), a step essential for releasing TCR⁺ microvesicles that trigger antigen-specific APC telomere donation^6,16^. The T_sen_ model therefore provided a system to restore FAO and directly test its effect on synapse formation and subsequent telomere acquisition, propaedeutic to River release.

Restoring CPT1A by lentiviral expression reactivated FAO in T_sen_ (ED Fig. 1c) and normalised TCR clustering (and thus TCR⁺ microvesicle release^6^) to levels seen in early-stage CD27⁺CD28⁺CD4⁺ T cells (T_erl_; **Fig. 2a** vs ED Fig. 1b). FAO lowered ceramide levels (ED Fig. 1d), relieved ceramidase inhibition (ED Fig. 1e-f), and restored phosphatidylethanolamine (PE) synthesis (**Fig. 2b, top**) — a lipid essential for synapse stability^17, 18^. Supplementing PE alone rescued TCR clustering, whereas sphingosine or S1P did not (**Fig. 2b**); conversely, ceramidase inhibition ^19^ abolished the CPT1A effect, directly linking FAO to synapse formation via the ceramidase–PE axis (Extended Data Fig. 1g).

**Figure 2:**
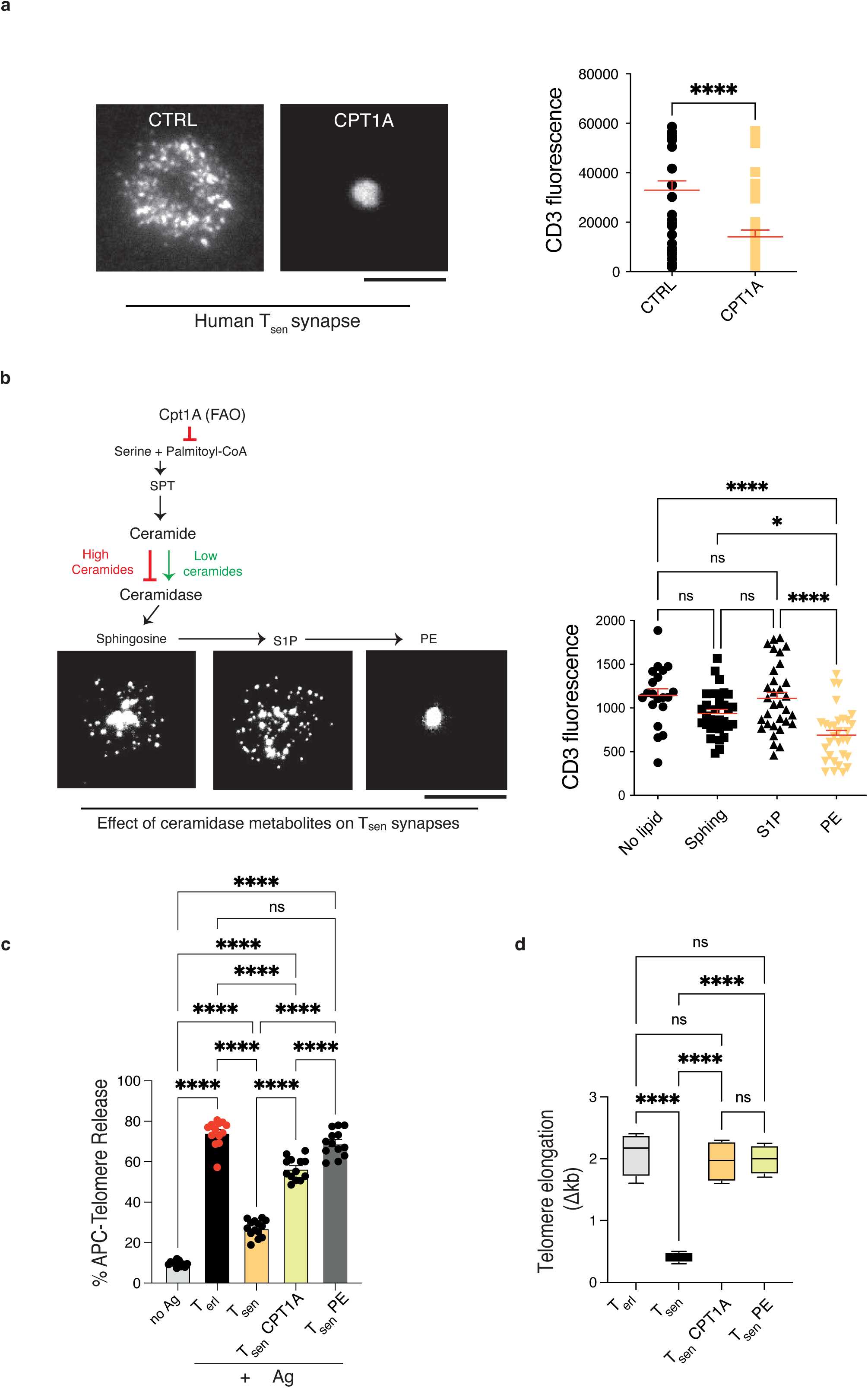
Propaedeutics to River formation. (**a**) Synapse-formation on planar bilayers of T_sen_ transduced with CTRL or CPT1A-expressing vectors showing that CPT1A-driven FAO restores TCR centralization at the synapse. Note that functional synapses have centrally-accumulated TCRs with lower CD3 fluorescence. Representative data (left) and quantification (right) of *n* = 35 synapses per group (*n* = 4 donors). Scale bars, 10 μM. (**b**) Ceramidase links FAO to the immune synapse. FAO activates ceramidase by blocking ceramide synthesis that in turn generates PE needed to centralize TCRs at the synapse. In contrast, short (20 min) treatment of T_sen_ with the PE precursors sphingosine or sphingosine-1 phosphate (S1P) did not centralize TCRs. Data are from *n* = 35 synapses per group (4 donors). Scale bar, 10 μM (**c**) Quantification of telomere release 24h after antigen-pulsed APC activation on TCR-coated bilayers. Data pooled from 4 donors (T_erl_, T_sen_, T_sen_ PE) or 2 donors (T_sen_ CPT1A). T_sen_ synapse competence was restored either by adding PE for 20 min or rescuing CPT1A expression by lenti-vectors for 10 days prior to synapse formation on bilayers. Antigen free APCs are shown, as control. (**d**) Telomere elongation by qPCR in metabolic competent T cells obtained by CPT1A expression or PE supplementation prior to conjugation with autologous antigen-pulsed APCs. T cells were purified from conjugates and analyzed 48 hours after synaptic activation (*n* = 4 donors). In (**a**) Mann-Whitney test and one-way Anova with Bonferroni post-correction for multiple comparisons in (**b**-**d**). *P<0.05, ****P < 0.0001. Error bars indicate s.e.m.

To model antigen-specific telomere transfer, we used planar lipid bilayers coated with anti-TCR antibodies to trigger synaptic TCR clustering and deposit TCR⁺ vesicles, as described^6^. After T-cell removal, only TCR microvesicles are maintained on the bilayer^6,16^, at which point TelC-labelled APCs —loaded or not with antigen mix (CMV, EBV, influenza, throughout)— were added. Antigen-loaded APCs released telomere vesicles in response to TCR⁺ vesicles, but only when FAO–ceramidase signaling had been restored in T_sen_, similar to non-senescent T_erl_ controls (**Fig. 2c**). Peptide-deprived APCs largely failed to release telomeres, confirming strict antigen dependency. FAO flux restoration also lengthened telomeres in recipient T cells (**Fig. 2d**), establishing FAO as a driver of telomere transfer at the immune synapse.

With the telomere mechanism acquisition defined, we next examined how Rivers formed from telomere-recipient T cells. We first applied DIA proteomics ^20^, comparing human Rivers with APC telomere vesicles. This analysis revealed marked depletion of glycolytic enzymes such as GAPDH ^21^, alongside enrichment for stem-related pathway proteins (**Fig. 3a**). Gene set enrichment analysis confirmed a strong bias towards stemness-associated pathways (Supplementary Information 4). This GAPDH^^low^, stem-factor–enriched signature was also detected in Rivers isolated directly from human serum using TelC–biotin pull-down followed by DDA proteomics (**Fig. 3b**), demonstrating that a hypoglycolytic, stemness-enriched profile is a conserved feature of human Rivers *in vivo*.

**Figure 3:**
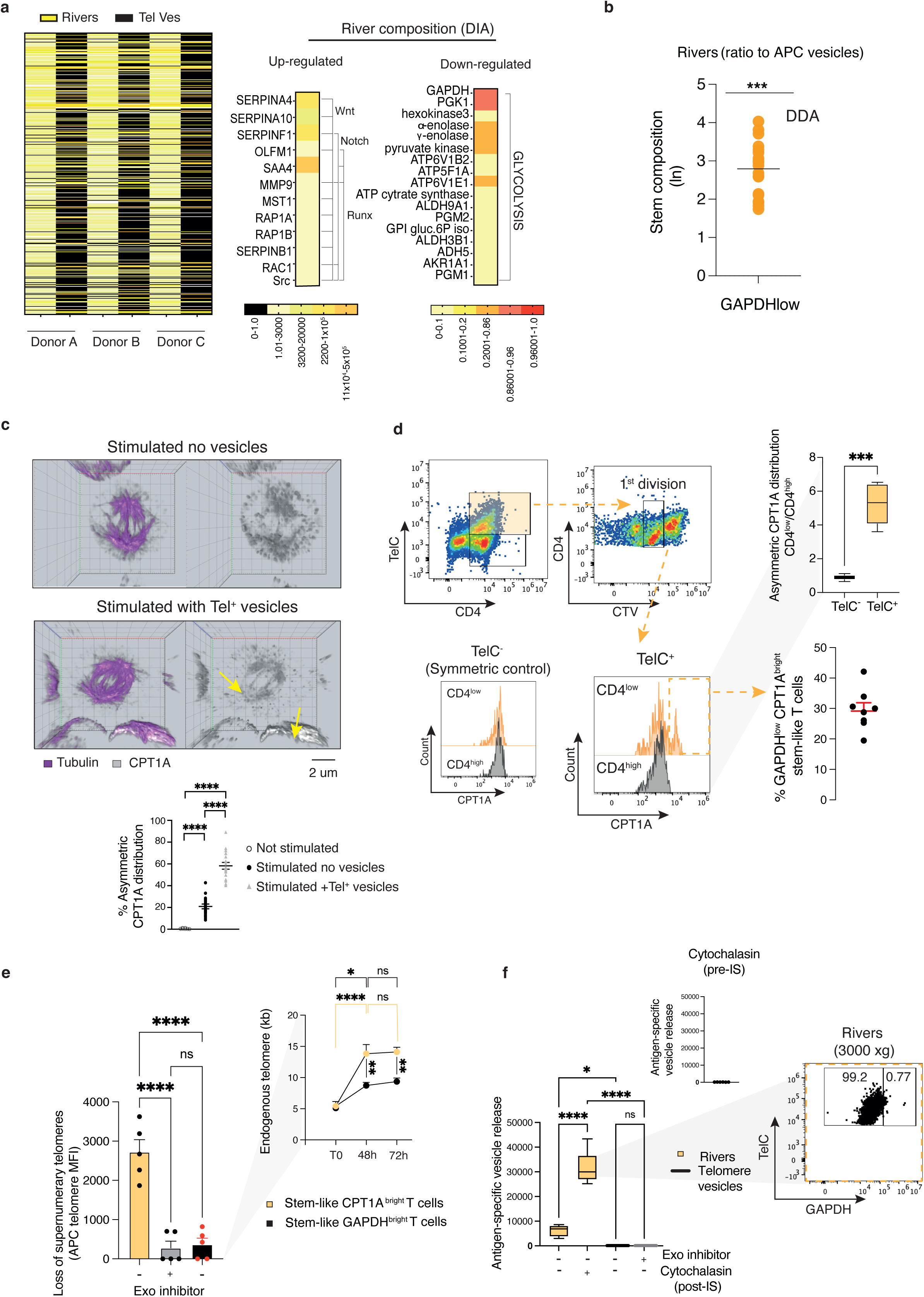
Generation of Rivers. (**a**) Proteomic comparison by DIA. Telomere vesicles and circulating Rivers were analyzed by LC–MS/MS with the data-independent acquisition (DIA) approach. Human APCs from each donor were stimulated with ionomycin overnight to induce vesicle release, and matched serum was collected in parallel. Heatmaps show differential abundance of glycolytic versus stem-related proteins (*n* = 3 donors per group). (**b**) Proteomics by DDA. Telomere-bound proteins were purified by TelC-biotin immunoprecipitation among and analyzed by LC–MS/MS with data-dependent acquisition (DDA). Rivers displayed enrichment of stemness factors and loss of GAPDH relative to APC telomere vesicles (*n* = 8 donors). (**c**) AD-like CD4^+^ T cell behavior. Representative IF–FISH images (top) and quantification from pooled microscopy (bottom; 17 images, n = 3 donors) showing asymmetric CPT1A distribution in T_erl_ cells after stimulation with anti-CD3/CD28 for 48 h, with or without telomere vesicles. Cytokinesis was blocked at 44 h; CPT1A asymmetry was assessed 4 hours later. Scale bar, 2 µm. Images were z-stacked and analyzed by confocal microscope. Scale bar, 2 μm (**d**) Flow cytometry of AD-like cells. Representative gating (*n* = 9 donors) showing asymmetric CPT1A distribution in telomere recipient (Tel⁺) versus non-recipient (Tel⁻) CD4⁺ T cells 48 h after conjugation with TelC-labeled APCs. Cells were tracked with CTV dyes, followed by flow-FISH (TelC) and CPT1A staining. GAPDH staining confirmed differential segregation in daughter cells. (**e**) Spontaneous loss of transferred APC telomeres. CPT1A^^bright^ versus GAPDH^^bright^ stem-like CD4⁺ T cells were analyzed 48–72 h after APC conjugation, with or without GW4869 (10 µM). Endogenous T-cell telomeres were elongated and remained stable, whereas transferred APC telomeres were selectively lost, indicating selective disposal of supernumerary APC telomeres (*n* = 5 donors). Telomere loss was calculated as ΔTelC fluorescence (48–72 h). (**f**) Release of Rivers by post-synaptic CD4⁺ T cells. After 24 h of APC–T cell co-culture, CD4⁺ T cells were freed by negative selection and cultured in isolation for 48 h. Supernatants were then analyzed for released telomeric particles. Rivers were defined as PKH67⁺ TelC⁺ GAPDH⁻, whereas APC vesicles were PKH67⁺ TelC⁺ GAPDH⁺. Rivers incorporated stemness factors (WNT5A, RUNX2; **Extended data fig. 2**). Pre-treatment with GW4869 or cytochalasin D blocked River release; by contrast cytochalasin D promoted River production when added 44 h after initial activation. Data are from (*n* = 6 donors). One-way Anova with Bonferroni post-correction for multiple comparisons in (**c**, **d**, **f left**) and paired Student’s t-test in (**b, f right**) were used. *P<0.05, ***P<0.001, ****P < 0.0001. Error bars indicate s.e.m.

We then asked how telomere-recipient T cells might modify River telomere cargo. Because FAO promotes asymmetric division (AD) ^22^ — a process in which daughter T cells inherit distinct metabolic and signaling components that specify fate ^23,24, 25^ — we hypothesized that telomere-recipient T cells might use an AD-like segregation mechanism to generate Rivers. In T_erl_ activated in the presence of purified telomere vesicles, CPT1A clusters were visible in mitotic cells, consistent with asymmetric distribution ^26^ (**Fig. 3c**). Using established AD criteria (asymmetric CD4 segregation with CTV dilution^23^) together with telomer transfer assays^6^, we identified a population of CD28⁺CD45RA⁺CD62L⁺CD95⁺ stem-like CD4⁺ T cells within the first division of APC telomere recipients that retained high CPT1A but reduced GAPDH expression (**Fig. 3d**; Supplementary Information 5). No equivalent population was observed in telomere-negative controls, and AD arrest via anti-LFA1^23^ antibodies disrupted GAPDH/CPT1A segregation (Supplementary Information 5).

Flow-FISH analysis revealed that while all stem-like (telomere recipient) T cell subsets elongated their endogenous telomeres, only CPT1A^bright T cells exhibited a reduction in supernumerary telomeres, consistent with their release into the supernatant (**Fig. 3e**). This reduction was abolished by pre-treatment with the exosome inhibitor GW4869^27^, indicating a vesicle secretion pathway. Telomere release did not compromise endogenous telomere elongation, suggesting that Rivers recycle surplus APC-donated telomeres without depleting T cell reserves.

To confirm River generation, we repeated antigen-specific T cell activation with autologous APCs, mechanically dissociated conjugates, and cultured isolated T cells for 48 hours without further stimulation. Rivers were detected mostly after antigen-specific activation and were enhanced by cytochalasin D at 44 hours, which halts further T cell division (**Fig. 3f**). By contrast, pre-activation cytochalasin D blocked River release entirely, showing that at least one round of T cell division is required for River production (**Fig. 3f, top right**). Sequential centrifugation at 3000 xg of T cell supernatants yielded GAPDH^^low^ telomeric material in large NET-like structures ^28^ (**Fig. 3f** and ED Fig. 2), physically distinct from the smaller GAPDH^^high^ telomere vesicles of APC origin (Supplementary information 6). Co-localisation and flow cytometry confirmed minimal, if any, GAPDH–telomere recruitment in Rivers, with a stem-factor profile closely matching that of circulating Rivers (ED Fig 2). Subset analysis identified stem-like CD4⁺ T cells as the dominant River producers, followed by central memory cells, whereas naïve, effector memory, and EMRA subsets released little or none; absence of synaptic APC activation also abolished T cell River formation (ED Fig. 3). Blocking telomere fusion in recipient T cells via siRAD51 knockdown in donor APC vesicles^6^ markedly increased River release (ED Fig. 4), further indicating that Rivers represent ‘spare’ APC telomeres that fail to integrate in the T cells after transfer.

**Figure 4:**
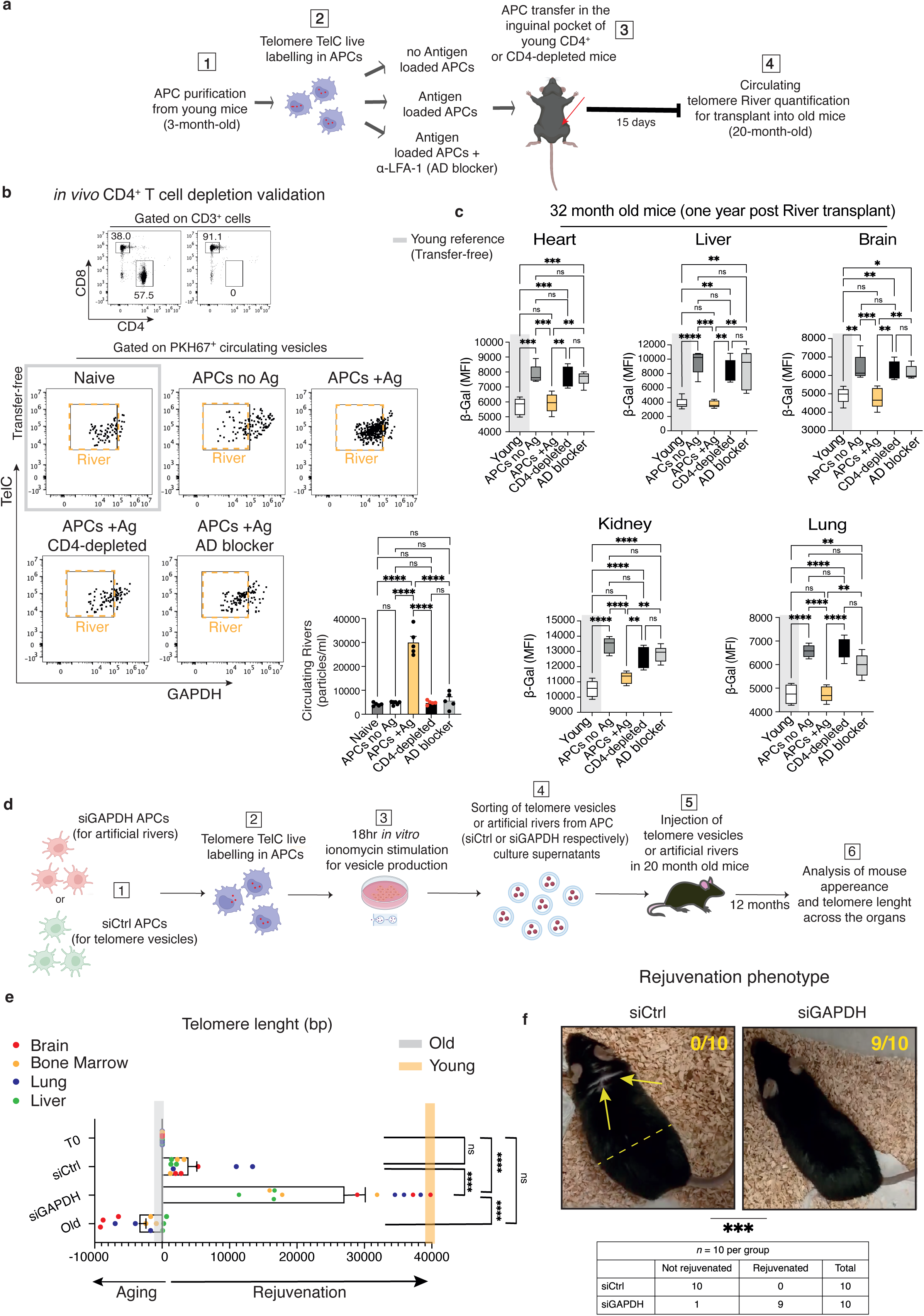
Systemic rejuvenation by telomere Rivers. (**a**) Experimental design for *in vivo* River generation with or without CD4⁺ T cell depletion. CD4-proficient or CD4-depleted recipient mice were adoptively transferred with telomere-labelled APCs, pre-loaded or not with Fluad vaccine as antigen source, with or without anti-LFA-1α antibody. Serum was collected 15 days later for River collection. Three months mice were used. (**b**) Representative FACS plots showing efficiency of CD4⁺ T cell depletion (top) and detection of telomere Rivers in each experimental group (bottom). Data are from (*n* = 5 mice per group). (**c**) Rejuvenating effects of telomere Rivers. Rivers were isolated from mouse serum as in **a** by FAVS sorting (PKH67⁺TelC⁺GAPDH⁻) and transplanted subcutaneously into 20-month-old congenic recipients. Animals were bred normally and assessed 12 months later. Flow cytometry of different organs demonstrated reduced senescence markers (β-Gal activity) and increased telomere length. For p16, IL-6, sestrin2-p-p38 (sMAC)), **Extended** Fig. 10) Data are from (*n* = 5 mice per group). MFI, mean fluorescence intensity. Young (3-month-old) mice are shown, as control. (**d**) Artificial River generation, experimental design. APCs were purified from total splenocytes of 3-month-old mice and transfected with siRNAs to silence GAPDH or with control siRNA. APC telomeres were then labelled with TelC probe and ionomycin stimulated *in vitro* to produce vesicles. Telomere vesicles (siCtrl vesicles) or artificial Rivers (siGAPDH vesicles) were sorted and injected into 20-month-old mice to test their rejuvenating effect. (**e**) Elongation of tissue telomeres by artificial Rivers. Telomere length was evaluated by Flow-FISH in multiple target tissues from the same 20-month-old mice as in **d**. Results are shown as absolute base-pairs (bp). Data are pooled from (*n* = 12 organs per group) and expressed as ΔTL relative to the old baseline at the start of the experiment (T0; grey bar), thereby distinguishing anti-aging effects (maintenance of old baseline; left) from rejuvenating effects (progression toward young reference values; right). siCTRL telomere vesicles largely delayed further telomere attrition without substantial rejuvenation, whereas artificial Rivers (siGAPDH) induced marked telomere elongation toward young reference values. Young (3-month-old) reference telomere lengths are indicated in yellow. In parallel, untreated old mice were aged normally to derive telomere attrition trajectories. (**f**) Appearance of representative mice injected with telomere vesicles (siCtrl vesicles) or with artificial telomere Rivers (siGAPDH vesicles). Note fur and body lean differences (*n* = 10 mice per group). One-way Anova for repeated measures with Bonferroni post-test correction (**b**-**e**). **P < 0.01, ***P < 0.001, ****P < 0.0001. Error bars indicate s.e.m.

APC telomere immunoprecipitation confirmed that GAPDH⁻ Rivers generated by T cells bound NOTCH1, β-catenin, and RUNX2 ^29–31^ upon recycling, whereas APC vesicles retained GAPDH and lacked stemness factors (ED Fig. 5). Donor-matched single-particle flow cytometry showed overlapping profiles between Rivers generated *in vitro* and natural Rivers from human plasma (ED Fig. 6). Silencing stemness factors in recipient T cells by siRNAs or genetic CRISPR abolished their detection in the Rivers, confirming T-cell origin; and dual-labelling returned River membranes derived from both T cells and APCs, whereas telomeric DNA was exclusively from APCs (ED Fig. 7). By contrast, CD8⁺ T cells failed to generate Rivers (ED Fig. 8), highlighting the selective capacity of CD4⁺ T cells to propagate this novel rejuvenating programme.

**Figure 5:**
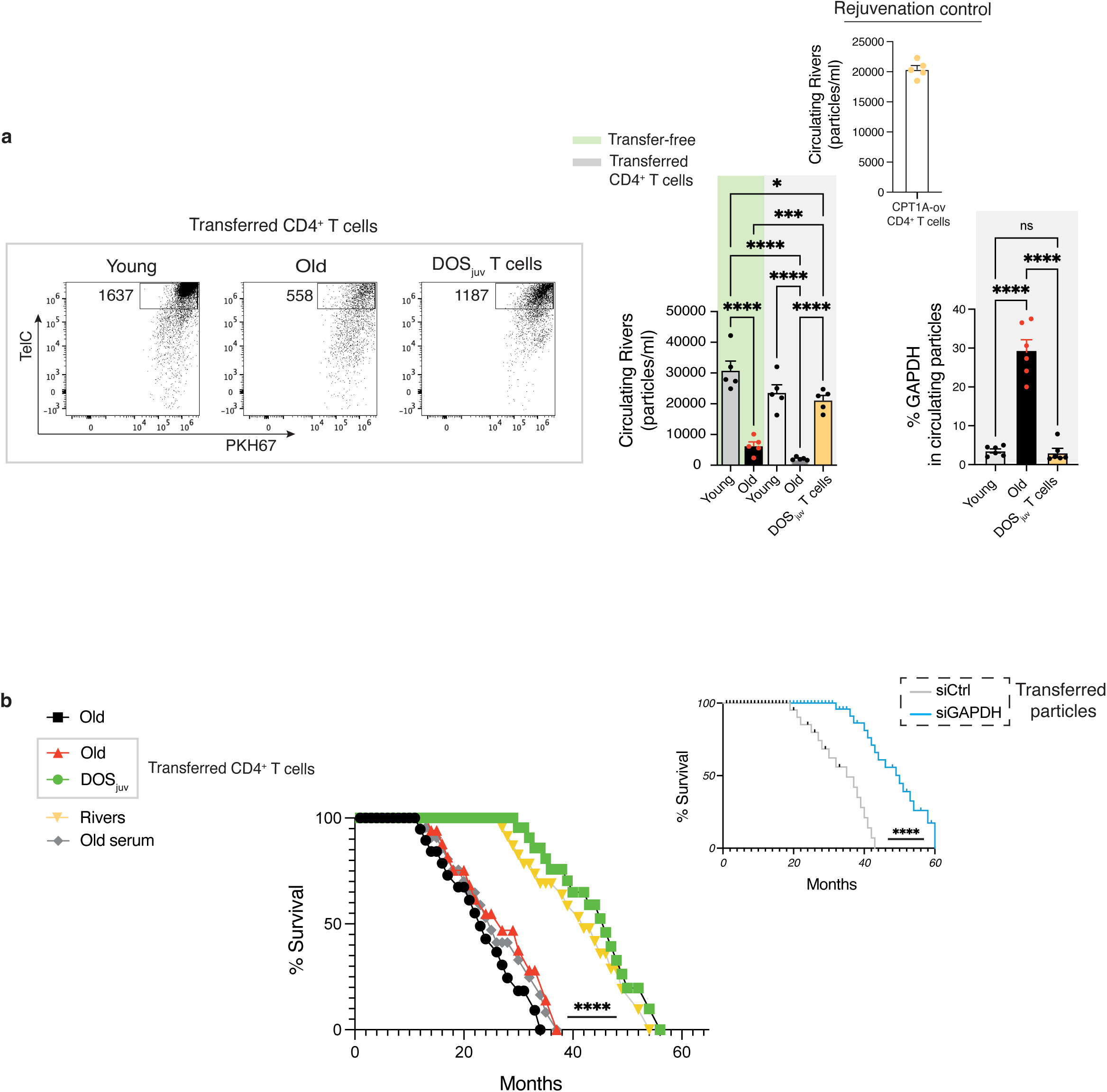
T cells extend lifespan via telomere Rivers. (**a**) CD4⁺ T cells were purified from the spleen of 20-month-old mice, treated or not with DOS for 4 h, and adoptively transferred into age-matched recipients. In parallel, young (3-month-old) and untreated old CD4⁺ T cells were also transferred. Some old CD4⁺ T cells were transduced with CPT1A vectors as an additional rejuvenation control. Recipient animals were vaccinated with Fluad vaccine 18 hours later and presence of circulating Rivers was studied 15 days after adoptive transfer. Representative FACS plots (left) and pooled quantifications (right) are shown. Transfer-free vaccinated animals were included as background controls. Transfer of rejuvenated (DOS or CPT1A) or young T cells strongly restored River production, whereas old T cells alone did not. Telomere Rivers were identified as PKH67⁺TelC⁺ vesicles. Note that flow cytometry confirmed GAPDH depletion in Rivers derived from young or rejuvenated serum, whereas old serum contained fewer telomeres with residual GAPDH contamination, consistent with disrupted River formation during aging (*n* = 5 mice per group for circulating Rivers; *n* = 6 per group for GAPDH staining). (**b**) Lifespan extension by telomere Rivers. Rivers as above (from DOS-rejuvenated T cell recipients) were FAVS-sorted and transplanted into age-matched old mice (∼5,000 particles/animal, single dose; *n* = 10). Parallel cohorts received artificial Rivers (siGAPDH vesicles, *n* = 10; siCTRL vesicles, *n* = 10), DOS rejuvenated CD4⁺ T cells (*n* = 8), old CD4⁺ T cells (*n* = 8), or untreated old serum (*n* = 8). Recipient animals were bred under standard conditions and monitored until natural death. Transplant-free old animals (20-month-old, *n* = 26) were followed in parallel to establish baseline survival. One-way Anova with Bonferroni post-correction for multiple comparisons in (**a**) and Cox regression analysis in (**b**). **P < 0.01, ***P < 0.001, ****P < 0.0001. Error bars indicate s.e.m.

Having established the cellular requirements and composition of Rivers, we next examined their generation *in vivo*. To test the requirement for cognate CD4⁺ T cell activation, we transferred fluorescently telomere–labelled APCs into young mice in CD4 proficient or deficient recipients (**Fig. 4a**). APCs were either pulsed with defined influenza antigens (adjuvanted Fluad vaccine) or left unpulsed. Additional groups received antigen-pulsed APCs pretreated with anti-LFA1 (to block asymmetric division ^23, 32^) or were CD4⁺ T-cell–depleted before APC transfer. Antigen-pulsed APCs triggered robust telomere River release *in vivo*, whereas unpulsed APCs, anti-LFA1–treated APCs, or CD4 depletion abolished River production, to background levels observed in naïve animals (**Fig. 4b**). Similar results were obtained using dendritic cells pulsed with a single OVA antigen in OT-II recipients, confirming dendritic cells as the main telomere donors among APCs (ED Fig. 9a); or with non-adjuvanted influenza vaccine–pulsed APCs demonstrating that River formation depends on antigen recognition rather than adjuvant formulation (ED Fig. 9b). Thus, CD4⁺ T cells are the source of telomere Rivers *in vivo*.

We next asked whether Rivers could reverse age-associated phenotypes. Circulating Rivers were sorted from equal serum volumes of young donors (∼140 μL from 250 μL whole blood), preserving natural abundance differences between conditions, using PKH67⁺ lipid dye, TelC⁺ telomeric DNA, and GAPDH⁻ status as sorting parameters. These Rivers were then transferred into aged recipients (20 months old). One year later (32 months), River-treated mice showed marked rejuvenation in brain, liver, lung, kidney, and heart; senescence-associated β-galactosidase activity ^33,34^ was reduced (**Fig. 4c**), sMAC (sestrin2^+^ p-p38^+^) formation diminished, and p16^INK4a and IL-6 expression decreased (ED Fig. 10). β-galactosidase positivity was confined to sMAC⁺ cells, indicating these markers identify the same senescent subset (Supplementary Information 7). Rivers from unpulsed APC transfers, CD4-depleted hosts, or AD-blocker groups failed to rejuvenate tissues, confirming the requirement for antigen-driven, cognate APC-CD4⁺ T cell interactions for River production *in vivo*. Rejuvenated tissues also showed reactivation of stem-related signaling, with increased NOTCH expression (ED Fig. 11a) and measurable telomere elongation across multiple organs (ED Fig. 11b).

To probe the role of GAPDH, we bypassed T cells by silencing GAPDH in APCs before vesicle release (**Fig. 4d**). Surprisingly, these GAPDH-depleted vesicles resembled natural Rivers — lacking glycolytic enzymes and containing stemness factors (ED Fig. 12) —perhaps due to competitive loading for stem factor vesicle content with GAPDH. When injected into old recipients, these “artificial” Rivers drove greater telomere elongation and tissue rejuvenation than natural Rivers or telomere vesicles (**Fig. 4e** vs ED Fig. 11b) and improved animal rejuvenated appearance (**Fig. 4f**). By contrast, GAPDH⁺ telomere vesicles from control APCs failed to restore telomere length or youth phenotype.

To assess whether CD4⁺ T cell–mediated rejuvenation also extended lifespan ^35^, we adoptively transferred young, old, or rejuvenated CD4⁺ T cells — this latter pre-treated with sMAC disruptors ^5,36,37^ (DOS - a specific T cell rejuvenating drug that activates FAO^37^) — into aged recipients, followed by influenza vaccination, to trigger antigen-driven River release (**Fig. 5a**). River production was restored in old animals receiving rejuvenated T cells, matching levels in young, immunised mice. Similar restoration occurred when T cells were rejuvenated by lentiviral CPT1A overexpression prior to transfer (**Fig. 5a**). In contrast, old T cells transferred into old recipients failed to restore River production. Rivers from young and rejuvenated T-cell recipients were largely GAPDH⁻, whereas those from old-to-old transfers retained GAPDH and were fewer in number (**Fig. 5a, right**).

Isolated Rivers (∼5.0 × 10³ particles) from these rejuvenated recipients were then transferred into old animals that did not receive T cells. River transplants extended median lifespan by ∼17 months and increased maximum lifespan, with several mice living to ∼58 months (**Fig. 5b**). Comparable effects were achieved with DOS-rejuvenated T cells or artificial siGAPDH Rivers, indicating that a single intervention can initiate a durable (and transferable) rejuvenation programme *in vivo*. Old serum containing predominantly GAPDH⁺ telomere vesicles, or young siCtrl GAPDH⁺ telomere vesicles, produced little or no effect (**Fig. 5b**). Thus, CD4⁺ T cells propagate youth- and lifespan-extending signals across individuals via telomere Rivers.

## Discussion

We identify telomere Rivers as an immune-derived route to systemic rejuvenation, a particle mediated process capable of transferring youth between organisms. Unlike telomerase, which acts within individual cells^36^, Rivers deliver telomeric DNA and stemness factors systemically, reversing senescence across multiple tissues. These results expand the role of T cell immunity well beyond canonical control of infections or cancers. In this respect, telomere Rivers resemble an immune-driven quorum-sensing pathway^38^, whereby CD4⁺ T cells coordinate systemic rejuvenation to sustain community healthspan – an unprecedent form of ‘herd’ rejuvenation. This is different from engineered CAR T cells that eliminate senescent cells^39^, as it acts via a transplantable element to spread reprogramming and rejuvenating signals across organisms.

The idea that circulating factors influence ageing has a long history, most notably in parabiosis ^1,2^, but reproducibility and mechanism have remained elusive. Whereas plasma exchange largely dilutes SASP factors, and soluble molecules^40^ act in organ restricted contexts, Rivers are the first discrete biological particles exerting systemic rejuvenation without reliance on blood dilution or organ-limited effects.

Rivers differ fundamentally from previously proposed factors. They are structured telomeric vesicle networks, generated by CD4⁺ T cells after telomere acquisition from APCs. Selective exclusion of glycolytic enzymes and enrichment in stemness pathways confer Rivers a unique molecular signature distinct from their APC-derived telomere precursors^6^.

Functionally, Rivers reverse senescence, extend lifespan, and act independently of endogenous telomerase, potentially buffering against telomere attrition in telomerase-deficient states^41^. Their production requires antigen-driven activation, FAO-dependent AD-like behavior of CD4^+^ T cells, and GAPDH exclusion – underscoring tight physiological regulation. The failure of APC telomere vesicles lacking stemness factors to rejuvenate recipients, compared with the robust systemic effects of Rivers, underscores stemness cargo as key determinant of rejuvenation. Correspondingly, silencing GAPDH in APCs generated stem enriched “Artificial Rivers” with strong rejuvenation potential.

By tracing their origin, composition, and systemic effects, we now define the first immune-regulated and transplantable programme of youth. The evolutionary conservation of Rivers across species—and even kingdoms—opens the prospect of engineering youth-promoting signals transferable between organisms, providing a conceptual framework for future immune-driven rejuvenation transplants. Youth transplant for community safeguard may thus represent a major yet overlooked function of T cells.

## Supporting information

Supplementary information; Lanna et al 2025

## Methods

### Ethics

All human and animal studies were performed in compliance with relevant ethical regulations. Human specimens were obtained with approval from the Tuscany Life Sciences Ethical Committee or the University of Oxford. Informed consent was obtained from all participants in accordance with the Declaration of Helsinki, and no financial compensation was provided. For mouse studies, protocols were approved by the Italian Ministry of Health. Mice were housed under controlled circadian rhythm (12 h light/12 h dark), constant temperature (25 °C), and humidity (40–60%). For human studies, age (20–60 years) and sex were evenly distributed across experimental groups.

### Human cell isolation and culture

Peripheral blood mononuclear cells (PBMCs) were isolated from healthy donor blood using Ficoll density gradient centrifugation (GE Healthcare, 17-1440-03). CD4⁺ T cells were purified by positive selection using CD4 MicroBeads (Miltenyi, 130-045-101). Senescent CD27⁻CD28⁻CD4⁺ T cells (Tsen) were obtained by negative selection with the CD4⁺ T Cell Isolation Kit (Miltenyi, 130-096-533), followed by depletion with CD27 and CD28 MicroBeads (Miltenyi). Early-differentiated CD27⁺CD28⁺CD4⁺ T cells (Terl) were isolated as non-senescent controls. CD3⁻ antigen-presenting cells (APCs) were obtained by CD3 depletion (Miltenyi, 130-050-101). All primary cells were cultured in RPMI-1640 (Sigma-Aldrich, R2405) supplemented with 10% fetal bovine serum (Sigma, F9665) and 1% penicillin-streptomycin (Lonza, DE17-602E), and maintained at 37 °C with 5% CO₂.

### Synapse planar bilayers

T_sen_ CD4⁺ T cells were activated for 48 h with anti-CD3 antibody and IL-2, followed by transduction with either control or CPT1A-overexpressing vectors (MOI = 10). After 10 d, cells were rested for 18 h at 37 °C before loading onto planar lipid bilayers prepared as previously described. In some experiments, CPT1A-transduced cells were pre-treated for 2 h with the ceramidase inhibitor Ceranib-1 (12 µM; Tocris). Lipid supplementation included phosphatidylethanolamine (PE, 2 µM, 20 min pre-treatment; Sigma), sphingosine (2 µM; Avanti Lipids), and sphingosine-1-phosphate (2 µM; Cambridge Bioscience). Synapse formation was assessed after 20 min stimulation by fixation in paraformaldehyde (10 min) and imaging with a 150× TIRF objective (Olympus). CD3 fluorescence was quantified using ImageJ (v2.1). Individual synapse experiments were recorded simultaneously to ensure identical raw fluorescence acquisition. For APC–telomere extraction quantification, after 20 min activation on bilayers, T cells were removed with cold PBS, leaving TCR microvesicles retained on the bilayer. Telomere-labelled (TelC) APCs loaded with or without CMV, EBV, and influenza lysates (Zeptometrix) were then added onto these TCR-microvesicle–enriched bilayers, and the percentage of APCs releasing telomeres was quantified 24 h later. Antigen-specific release was determined by subtraction of background values obtained from antigen-free APCs.

### Western blotting

Purified T_erl_ (CD27⁺CD28⁺CD4⁺), T_int_ (CD27⁻CD28⁺CD4⁺), and T_sen_ (CD27⁻CD28⁻CD4⁺) T cells were lysed in RIPA buffer (50 μL per 10⁶ cells; Sigma, R0278) with protease and phosphatase inhibitors on ice for 30 min, followed by centrifugation (15,871 × g, 20 min, 4 °C). Protein extracts were denatured in Laemmli buffer with β-mercaptoethanol, separated by SDS-PAGE, and transferred to nitrocellulose membranes. After blocking in 5% non-fat milk, membranes were incubated overnight at 4 °C with primary antibodies against CPT1A (Cell Signaling, 12252, 1:1000), SPT (Sigma, AV46721, 1:1000), or AMPK-α (Cell Signaling, 2532, 1:1000).

Secondary HRP-conjugated antibodies (Cell Signaling, 7074P2; Sigma, 12-349; 1:5000) were applied, and signals detected using ECL (Thermo, A38554) and imaged on Chemidoc (Thermo iBright1000).

### Metabolic assays

Fatty acid oxidation (FAO) was measured using an ELISA-based kit (Abcam, ab217602). Tsen cells transduced with CPT1A or control vectors were treated with the SPT inhibitor myriocin (10 µM, Sigma) in the presence or absence of BSA-palmitate (100 µM), followed by ceramide quantification. Ceramidase activity was assayed after ASAH2 immunoprecipitation (Abcam, ab113369) from 10⁷ T_sen_. Cell lysates were prepared in RIPA buffer, either directly ex vivo or 96 h after mock or CPTIA vector transduction. Enzymatic conversion of ceramide to sphingosine was measured by o-phthaldialdehyde fluorescence (excitation 340 nm, emission 450 nm). Exogenous C12-ceramides (Avanti Polar Lipids; 50–2000 µM) were added in some assays. Ceramide abundance was measured by ELISA (Sigma, MID 15B4). PE levels were quantified using Cys-fluorescent duramycin (Molecular Targeting Technology; MTTI, D-1002).

### Mass spectrometry assessment with Data Independent Acquisition (DIA)

Donor-matched telomere Rivers and vesicles were derived from sera or from ionomycin-activated APCs, respectively, and subjected to ultracentrifugation to enrich for sufficient material for proteomic studies. Pellets were then lysed with appropriate buffer (8M urea-Tris HCl; 20 min at room temperature). Next, pellets were loaded into 10K Millipore columns and centrifuged for 15 min at 14,000 xg, at 4 °C. Washing buffer was lysis buffer (3 washes, 15 min each at 14,000 xg). The 50 ul retention volume, containing the purified material, was eluted upon column inversion as this material is trapped in the 10K Millipore column matrix. Proteins were quantified by the BCA method and then analyzed with LC-MS/MS (Orbitrap nanoUPLC; Ceinge) coupled with the direct DIA approach (Spectronaut 18.5.231110.55695 software, Creative Proteomics, USA). A total of 1159 proteins were identified for natural Rivers and telomere vesicles. Intensive bioinformatics analyses were carried out to analyze those quantifiable proteins, including heatmaps.

Proteins of relative quantity were divided into two categories. Quantitative ratio over 1.2 was considered up-regulation while quantitative ratio less than 1/1.2 was considered down-regulation and this served as cut-off. For analysis, the fraction of background vesicles devoid of telomeres was subtracted. This fraction was derived from parallel flow-cytometry assessment with TelC probe and PKH67 dye from the same samples subjected to mass spectrometry.

For network-based pathway analysis, differentially abundant proteins identified by DIA proteomics (fold change >1.2 or <0.83; FDR <0.05) were subjected to multi-platform pathway enrichment. Enrichment analyses were performed using GSEA (MSigDB v7.5.1 Hallmark, KEGG, Reactome, and Gene Ontology Biological Process collections), g:Profiler (v0.2.3; with hierarchical filtering), and Enrichr (KEGG 2021, GO BP 2021, and BioPlanet 2019). Overlapping pathway annotations across platforms were consolidated into a unified enrichment table with FDR <0.05 as the significance threshold. For network reconstruction, significantly enriched proteins and their associated pathways were mapped to protein–protein interaction data from the STRING database (confidence score >0.7, experimental + curated interactions only).

The resulting interaction matrix was imported into Cytoscape (v3.9.1) for graph construction. Network topology was organized using a force-directed layout, with manual refinement for readability.

Nodes:

- Core pathway hubs (orange): canonical stemness-associated modules, including WNT, NOTCH, and RUNX1 signaling, which formed the central regulatory scaffold of the network.
- Stemness convergence module (green): clusters enriched for telomere maintenance, cellular multipotency, and mitochondrial non-oxidative resilience.
- Upregulated effector nodes (blue): individual proteins upregulated in Rivers (e.g., SERPINA4, SERPINA10, SERPINB1, OLFM1, SAA4), connected to stemness hubs via KEGG/GO overlap. Edges represent high-confidence regulatory associations inferred from STRING (confidence score >0.7), KEGG/GO curated pathways, and co-expression topology. Node degree and betweenness centrality were calculated in Cytoscape to identify hubs. Functional sub-networks were further clustered using the MCODE algorithm to highlight densely connected modules.

This integrated network framework captured hierarchical relationships between stemness-associated master regulators, downstream effector genes, and stress-resilience pathways, providing a systems-level map of the rejuvenation signature conferred by telomere Rivers.

### Mass spectrometry assessment with Data Dependent Acquisition (DDA)

APC vesicles or telomere Rivers were obtained as above described and incubated with TelC-biotinylated probes, followed by immunoprecipitation with streptavidin beads to isolate telomere complexes. Telomere bound proteins were then subjected to proteomics with data dependent analysis (DDA), suitable to identify purified protein extracts or complexes. Protein lysates were first digested on S-Trap cartridges, following the manufacturer’s protocol (Protifi). Sample were initially mixed with 5% SDS and 20 mM DTT, boiled, cooled to room temperature and then alkylated with 40 mM iodoacetamide in the dark for 30 minutes. Subsequently, a final concentration of 1.2% phosphoric acid and six volumes of buffer were added to the samples to allow the proteins to adsorb onto the S-trap cartridges. The protein solution was then loaded onto the S-Trap filter, centrifuged at 2000 rpm. The filter was then washed with binding buffer three times. Finally, digestion buffer containing trypsin (1:10 weight ratio) was added to the filter and digested. The peptide solution was then collected and freeze-dried for LC-MS/MS analysis. The peptide mixtures resulting from hydrolysis with trypsin were suspended in a volume of 0.2% formic acid until a concentration of 0.25 ug/ul was obtained. The mixtures were analyzed by LC-MS/MS analysis with an ORBITRAP EXPLORIS240 mass spectrometer (ThermoScientific, Waltham, MA) equipped with a Vanquish-Neo UHPLC system. The peptide mixture was first concentrated and desalted with a pre-column (PepMap™ Neo Trap Cartridge) then fractionated on a reversed phase C18 capillary column (Double nano Viper™ PepMap™ Neo 2 um C18 75 um X 150 mm) at a flow rate of 250 nl/min.

The peptide mixture was separated with a 77-minute chromatographic gradient using buffers A and B whose composition is reported below:

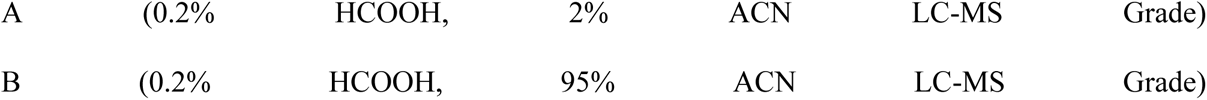

Peptide analysis was performed in “Data Dependent Acquisition” (DDA) mode, selecting the first 20 most intense ions (top 20) for each scan (range between 300 and 1800 m/z), with charge state >1.

Each sample was analyzed in technical triplicates. For peptide identification, the raw data obtained from the LC-MS/MS analysis were processed using the MaxQuant software, with the UniprotKB database considering the “Homo Sapiens” taxonomy. The requirements set for identification were the followings: Modifications: Oxidation (M); Gln-> pyro-Glu (N-Term Q); Carbamidomethyl (C); Enzyme: Trypsin; Max Missed cleaves: 2; Minimum number of peptides per ID: 3; Minimum number of unique peptides per ID: 1; FDR for protein identification: 0.01. Statistical analysis was performed using the Perseus platform, filtering for FDR<0.05. Several proteins belonging to stem related pathways were identified. GAPDH low identity of the immunoprecipitated telomere rivers was also confirmed in the DDA analysis itself, as compared to APC telomere vesicles.

### Generation of telomere Rivers

#### Natural Rivers

Natural telomere Rivers were generated in vivo by antigen-dependent CD4⁺ T cell–APC interactions. Three-month-old male C57BL/6 mice were used as recipients, with or without prior depletion of CD4⁺ T cells. CD4^+^ Depletion was achieved by two sequential intraperitoneal injections of anti-CD4 monoclonal antibody (InVivoPlus anti-mouse CD4, clone GK1.5; Bio X Cell, 300 µg on day 0 and 200 µg on day 2). For APC donors, spleens were harvested from naïve age-matched mice, and total splenocytes were subjected to negative selection with CD3ε MicroBeads (Miltenyi, 130-094-973) to remove T cells.

The CD3⁻ fraction was plated for 2 h to allow adherence, enriching for antigen-presenting cells (APCs). Adherent APCs were live-labelled with TelC-Cys probes (F1003; PNA Bio) using glass microbeads. APCs were then pulsed with the multivalent influenza vaccine Fluad (Seqirus; 1:50 of the human dose) for 3 h at 37 °C. Where indicated, cells were additionally treated with anti-LFA-1α blocking antibody (InVivoMAb anti-mouse CD11a, clone M17/4; Bio X Cell, 1 µg per 10⁶ cells) immediately prior to transfer.

Twenty million APCs were injected subcutaneously into the inguinal pocket of recipient mice. Control mice receiving antigen-free APCs were not vaccinated and served as naïve controls.

Serum (140 µl per animal) was collected on day 15 for analysis of circulating telomere Rivers. Rivers were pre-cleared at 3000×g and analyzed by flow cytometry using dual labelling with TelC probe and PKH67 dye. FAVS sorting was performed on a MoFlo Astrios EQ (Beckman Coulter; 5 lasers, 355–640 nm) using a gating strategy that excluded surface GAPDH⁺ events.

Sorting consistently achieved ∼97% isolation GAPDH-River particles. Identical serum input volumes were processed across experimental groups and were not artificially normalised post sorting, such that differences reflected natural River concentration exclusively. Alternatively, CD11c⁺ dendritic cells (DCs) were isolated from spleens of young C57BL/6 mice using the Pan-DC isolation kit (Miltenyi, 130-100-875). DCs were pulsed with OVA peptide (OT-II 323–339, 1 µg/ml; InvivoGen) for 3 h at 37 °C before transfer into OT-II recipient mice, with or without prior CD4⁺ T cell depletion. Formulation controls were performed using non-adjuvanted influenza Fluarix vaccine.

#### Artificial Rivers

Splenic antigen-presenting cells (APCs) were isolated from 3-month-old C57BL/6 mice and transfected with small interfering RNAs (siRNAs) to selectively silence GAPDH (sc-35449, Santa Cruz Biotechnology) or with a non-targeting control siRNA (sc-44236, Santa Cruz). Transfection was performed using Lipofectamine RNAiMAX (Thermo Fisher) following the manufacturer’s protocol, and efficiency was verified by qPCR and western blot analysis 48 hours post-transfection. APC telomeres were then live-labelled with TelCys probes (PNA Bio, F1003) under non-denaturing conditions. Cells were subsequently stimulated in vitro with ionomycin (0.5 µg/ml, 18 hours) to induce vesicle release. Supernatants were cleared by low-speed centrifugation (800 ×g, 10 min; 3,000 ×g, 1 hours, 4 °C), and vesicles were isolated by fluorescence-assisted vesicle sorting. Sorted vesicles were washed in PBS/0.1% BSA, quantified by Nanoparticle Tracking Analysis (NTA), and normalized to equal input across groups. Artificial Rivers or control vesicles were then transplanted subcutaneously into 20-month-old recipient mice. Animals were maintained under standard housing conditions for 12 months post-transplant.

### Adoptive transfer

CD4⁺ T cells were purified from total splenocytes of 20-month-old C57BL/6 mice using magnetic bead negative selection (Miltenyi). Cells were rejuvenated ex vivo by either: (i) treatment with the sMAC disruptor DOS (10 µM, 4 hours 37 °C) or (ii) Cpt1A overexpression by lentiviral transduction. Untreated old CD4⁺ T cells served as controls. In parallel, CD4⁺ T cells from young (3-month-old) mice were isolated and processed identically as reference controls. Rejuvenated CD4⁺ T cells, untreated old CD4⁺ T cells, or young CD4⁺ T cells were adoptively transferred into age-matched (20-month-old) recipient mice via intravenous injection. Recipient mice were immunized with Fluad vaccine (Seqirus; 1:50 human dose equivalent) at 18 hours post-transfer. Circulating telomere Rivers were assessed in serum samples 15 days later by flow cytometry using TelC probe (PNA Bio) and PKH67 membrane dye (Sigma). For baseline controls, young (3-month-old) and aged (20-month-old) mice not subjected to adoptive transfer were vaccinated in parallel.

### River detection by flow cytometry

Human APC telomeres were live-labelled with TelC-Cy5 probe (F1003, PNA Bio) as previously described and stimulated with ionomycin (0.5 µg/ml, 18 hours). Supernatants containing APC telomere vesicles were added to autologous purified CD4⁺ T cells, followed by activation with CD3/CD28 TransAct (130-111-160; Miltenyi) for 48 hours. Where indicated, T cell cultures were exposed to cytochalasin D (10 µM; Merck) 4 hours before supernatant collection. Supernatants were sequentially cleared at 800×g (10 min) to remove debris and at 3,000 ×g (1 hour, 4 °C) to pellet large vesicles, including River particles. To stabilize vesicles and prevent loss during washing, pellets were resuspended in 20% PEG-8000 and centrifuged again; the retained material was then permeabilized with Triton X-100 (0.001%, 10 min) and stained for GAPDH (5174; Cell Signaling), ENO1 (945502; BioLegend), PGK1 (68540; Cell Signaling), WNT5A (2392; Cell Signaling) or RUNX2 (12556; Cell Signaling). River membranes were labelled with PKH67 (Sigma) and analyzed by flow cytometry, defined as PKH67⁺ TelC⁺ GAPDH⁻ particles. These were enriched for stem-related enzymes. River particles are present in the larger vesicle fraction, likely due to the presence of NET-like structures. For comparison, telomere vesicles were derived from the same telomere-labelled APCs activated with ionomycin, but without T cell co-culture. Telomere vesicles were purified by ultracentrifugation (105,000 ×g) followed by FAVS, and were enriched for glycolytic enzymes but largely devoid of stem-related proteins. Alternatively, vesicles were assessed without selective enrichment by applying PEG-based precipitation directly to supernatants (3,000 ×g), enabling quantification of River particles among the total vesicle pool.

### River detection by flow cytometry

Human serum was obtained from healthy blood donors. Whole blood was layered onto Ficoll and centrifuged at 2,000 rpm for 20 min to separate plasma/serum fractions. Mouse serum was collected from peripheral blood following retro-orbital or submandibular bleeding and centrifuged at 2,000 rpm for 10 min.

In both cases, sera were first clarified at 800 ×g (10 min) to remove debris, followed by 3,000 ×g centrifugation to pellet large vesicles containing River particles. Pellets were directly stained with TelC probe (1:50 dilution, PNA Bio). To confirm telomeric specificity, staining was validated in parallel with anti-TZAP antibody, which recognizes the telomere-trimming factor TZAP, confirming probe binding to telomeric DNA directly ex vivo.

For plant xylem, orchid roots were collected, washed in PBS, and flushed with PBS using a syringe to obtain xylem fluid. Plant fluids were centrifuged at 800 ×g (10 min) to remove debris, followed by centrifugation at 3,000g. Pellets were stained with TelC probe and PKH67 dye as described for mammalian samples. In environmental modulation experiments, orchids were exposed to alternating 12 hour light / 12 hour dark cycles (two full cycles) or subjected to enforced dark–dark cycles (48 hour total). In the latter condition, plants were hydrated with or without fenofibrate (1 mM in 4.7% ethanol/PBS solution added to the soil). Xylem-derived Rivers were analyzed by flow cytometry at the end of treatment.

### River composition controls

To determine the origin of River membrane lipids, APCs or T cells were labelled or not with PKH67 before River induction, and membrane fluorescence among released Rivers was quantified by flow cytometry. To trace telomere origin, APCs and T cells were differentially labelled with TelC-Cy5 versus TelC-Cy3 probes, followed by River induction, enabling discrimination of telomeric sources in released particles.

To assess stem enzyme incorporation, CD4⁺ T cells were transfected with siRNAs targeting RUNX2, WNT5A, or NOTCH1. Stemness depletion among released Rivers was verified by flow cytometry after co-culture of these T cells with APC telomere vesicles and subsequent activation. To test whether Rivers incorporated excess telomeric material under conditions of impaired APC-mediated telomere elongation, APCs were transfected with siRNA against RAD51. RAD51-deficient APC vesicles, defective in telomere extension, were co-cultured with T cells. For all composition control experiments, Rivers were analyzed in the context of total vesicle release. Supernatants were subjected to PEG-assisted precipitation (20% PEG, centrifugation at 3,000 ×g), ensuring capture of all vesicle populations for comparative analysis. The resulting Rivers contained supernumerary telomeric content, consistent with increased recycling. To confirm River T cell stem factor origin, in some experiments CD4+ T cells were subjected to Runx2 KO by CRISPR technology (Origene; KN212884RB)

### River transplants

Telomere River particles were purified by FAVS sorting as described above and transplanted subcutaneously into 20-month-old C57BL/6 mice. Animals were housed under standard conditions, bred normally, and sacrificed 12 months later for assessment of systemic rejuvenation across multiple organs (liver, brain, kidney, heart, lung, and bone marrow). Tissues were analyzed for senescence and stress markers, including β-Gal (Cell Signaling #2372), p16 (Cell Signaling #29271), IL-6 (Cell Signaling #12912), sestrin2 (Abcam EPR18907), and phosphorylated p38 (Cell Signaling #4511). Telomere length was measured by flow-FISH as previously described2. In parallel, artificial Rivers were generated from APC-derived vesicles engineered by GAPDH silencing (siGAPDH) and purified by FAVS. Control vesicles (siCTRL) and artificial Rivers were transplanted subcutaneously into 20-month-old mice, which were then housed and bred normally for 12 months.

Animals were monitored longitudinally for phenotypic changes (fur quality, motor activity, posture, body weight, body composition). At sacrifice, telomere length (TL) was assessed in lymphoid and non-lymphoid tissues by flow-FISH. Age-matched untreated old mice (20-month-old) and young mice (3-month-old) served as references. Anti-ageing effects were defined as TL maintenance at the old baseline (no further attrition from the T0 reference), whereas rejuvenation was defined by a shift of TL distribution toward young reference values.

Untreated old animals were maintained in parallel to establish ageing trajectories.

### AD-like behaviour of River-producing T cells

Human CD27⁺CD28⁺ CD4⁺ T cells were pre-labelled with CellTrace Violet (CTV; Thermo Fisher) and conjugated with autologous telomere-labelled APCs. APCs were labelled with TelC-Cy3 probe (PNA Bio) and pulsed with an antigen mix (Hexyon, Sanofi, 1:100; Fluad, Seqirus, 1:100; PepTivator CMV pp65, Miltenyi, 1:50). T cell–APC co-cultures were maintained for 48 or 72 hours before analysis. For telomere length assessment, cells were subjected to flow-FISH using a second TelC probe with a distinct fluorochrome (TelC-Cy5). Absolute telomere length was determined with reference standards of known telomere length, as previously described, combined with intracellular detection of GAPDH (Cell Signaling #5174) and CPT1A (Abcam #ab128568). Because APC-derived telomeres carried a distinct fluorochrome, loss of their signal in the absence of endogenous T cell telomere shortening was interpreted as disposal of supernumerary telomeres not integrated into the T cell genome. In some experiments, CD4⁺ T cells were pretreated with the exosome release inhibitor GW4869 (10 µM, 1 h; Sigma-Aldrich) before APC conjugation. Additional controls included blockade of synapse integrity with anti-LFA-1α antibody (InVivoMAb, 1 µg/ml).

### Imaging

Samples were cyto-spun on glass slides for 5 min at 500 rpm, dried overnight at room temperature, and dehydrated by acetone–ethanol washing for 10 min each. Air-dried sections were stored at −80 °C for 4 hours to allow cell permeabilization prior to antibody stainings. Cervical lymph nodes were harvested from six-week-old (wild-type) C57BL/6J mice 7 days post re-challenge with ovalbumin and snap frozen using liquid nitrogen in CryoMoulds (VWR) containing OCT (Tissue Tek). Unchallenged (naïve) age-matched animals served as control. Tissue blocks were stored at −80 °C prior to fixation. Six-micron (6 μm) cryosections were analyzed by Immuno-FISH (IF-FISH) with telomere probes.

For telomere IF-FISH, samples were fixed in 2% paraformaldehyde for 20 min, washed in PBS, and treated 20 min with −20 °C ethanol. Blocking was performed for 1 h at room temperature in PBS-TT (8% BSA, 0.5% Tween-20 and 0.1% Triton X-100 in PBS). Primary stainings were performed overnight at 4 °C with antibodies to CD3 (1:1000; MCA463G; BIO-RAD), Tubulin (2144; Cell Signaling), CPT1A (1:100; 8F6AE9; Abcam), GAPDH (1:100; 5174; Cell Signaling). Secondary stainings were performed for 1 h at room temperature in the dark with donkey anti-Mouse IgG Alexa Fluor 647 (1:250; A32787; ThermoFisher) or goat anti-rabbit IgG Alexa Fluor 405 (1:250; A48254; ThermoFisher).

For IF-FISH, cross-linked sections (20 min in 4% paraformaldehyde) were subjected to dehydration in graded ethanol followed by hybridization with 40 pM PNA probe solution (TelC-Cy3; F1002; PNA Bio) in hybridization buffer (1 M Tris pH 7.2; magnesium chloride buffer; deionized formamide; blocking buffer, Roche; deionized water). DNA denaturation (10 min at 82 °C) was followed by 2-hour hybridization at room temperature. After washing in 70% formamide, saline sodium citrate (SSC) buffer and PBS, sections were mounted by Vectashield + DAPI (Vector) or Pro-Long Gold (Invitrogen). Imaging was performed using super-resolution ZEISS LSM 800 Airyscan (Zeiss) with Zen lite software version 2.3.

Sections were z-stacked and the integrated signals analyzed by ImageJ software (V2.1).

### Biochemical assessment of telomere Rivers

For biochemical assessment, human APCs were live-labelled with TelC-biotin probes (PNA Bio) and stimulated with ionomycin as described above, to derive telomere vesicles suitable for immunoprecipitation with streptavidin agarose beads (20347; ThermoFisher). The vesicles were then transferred, or not, to autologous human CD4⁺ T cells from the same donors. T cells were cultured in the presence of vesicles for 48 hours, with cytochalasin D added 4 hours prior to the end of the culture. Supernatants were collected, cleared by centrifugation at 800 ×g for 10 minutes, and subjected to immunoprecipitation in the presence of Triton X-100. The precipitated APC-telomeres from T cell cultures were directly analyzed by ELISA using antibodies against GAPDH (Cell Signaling #5174), Enolase (BioLegend #945502), PGK (Cell Signaling #68540), NOTCH1 (Cell Signaling #3608), Wnt5 (Cell Signaling #2392), and RUNX2 (Cell Signaling #12556). Results were recorded as absorbance at 450 nm on an ELISA microplate reader (AMR-100, Biolab).

### Life extension studies

Rivers were isolated 15 days after adoptive transfer of rejuvenated (DOS treated) CD4⁺ T cells into 20-month-old mice vaccinated with Fluad (twice, 18 hours and day 5). Blood (∼250 µl) yielded ∼140 µl serum, from which circulating Rivers were FAVS-sorted. Each transplant consisted of ∼5,000 River particles, delivered subcutaneously into age-matched old recipients not subjected to T cell transfer. Animals were then bred normally and monitored until natural death. Rivers were transplanted in each animal once. Parallel cohorts received artificial Rivers (siGAPDH telomere APC vesicles), systemic DOS, DOS treated CD4⁺ T cells, or Rivers derived from young T cells, each tested against appropriate controls (siCTRL telomere APC vesicles, untreated old serum, or untreated old T cells). Control mice received equal serum volumes (140 µl) from old donors, which contained mainly GAPDH⁺ telomere vesicles rather than Rivers.

### Statistical analysis

Statistical analyses were performed using GraphPad Prism v9. For human experiments, two-tailed paired Student’s t-tests were applied. For mouse experiments, unpaired t-tests or one-way ANOVA with Bonferroni correction were used as appropriate. Mann–Whitney tests were applied for imaging data. Kaplan–Meier survival curves were compared with Mantel–Cox analysis. P values <0.05 were considered significant.

## Data availability

The data generated or analyzed in this study are included in the manuscript, its Extended data and Supplementary information files. Mass spectrometry data are provided in the fig Share at submission. Source data are provided with this paper.

## Contributions

A.L. discovered the telomere Rivers and further conceived of, designed, performed, supervised and directed the study, analyzed and interpreted the data, provided fundings and laboratory infrastructures and wrote the manuscript. F.R. performed and analyzed experiments under supervision of A.L. S.V. performed bilayer construction and TIRF experiments with A.L. M. L. D. provided synaptic bilayer infrastructures.

## Acknowledgments.

This work was funded by Sentcell ltd. Bilayer synapse studies were supported by the Wellcome Trust 110229/Z/15/Z to A.L. The funders had no role in study design or decision to publish the manuscript. A.L. is a Honorary Professor of the University College London and the Chief Executive Officer of Sentcell ltd. MLD, SV were supported by the Wellcome Trust (100262Z/12/Z) and the Kennedy Trust for Rheumatology Research.

---

**Extended Data Fig. 1:**
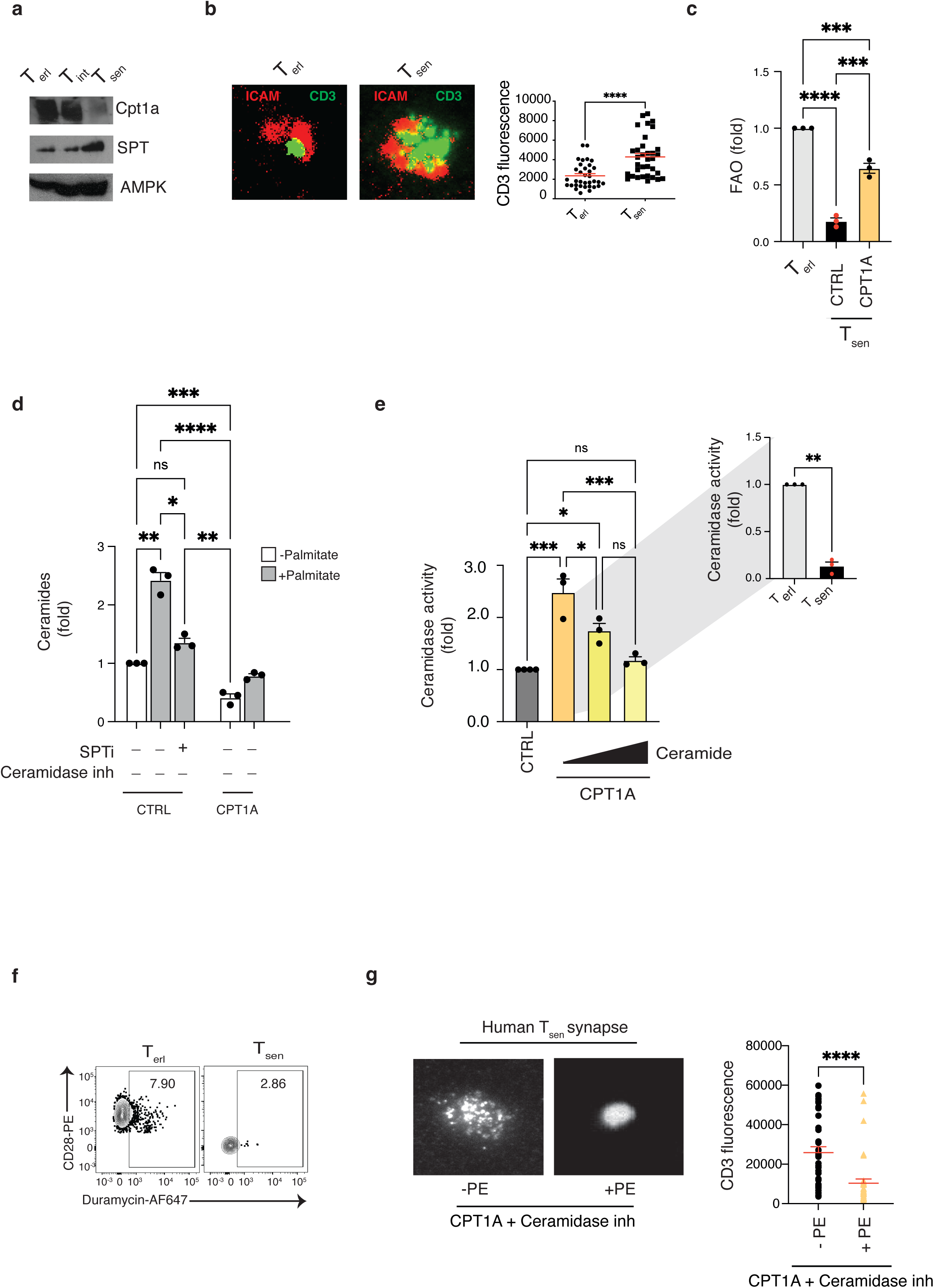
Ceramidase links FAO to the synapse. (**a**) Representative immunoblots of CPT1A and SPT proteins in human CD4^+^ T cells defined by relative CD27 and CD28 expression as T_erl_ (CD27^+^ CD28^+^), T_int_ (CD27^-^ CD28^+^) and T_sen_ (CD27^-^ CD28^-^). Note the loss of CPT1A and increased SPT, consistent with a metabolic switch to ceramides synthesis in senescent cells^14^. AMPK, loading control. SPT, serine- palmitoyltransferase. Representative of 4 separate donors. (**b**) Planar bilayers showing disrupted synapse formation in T_sen_ vs T_erl_. Representative images of 3 separate donors (**left**) and pooled data (**right**; *n* = 34 synapses). Purified cells were rested overnight before stimulation on supported lipid bilayers containing anti-CD3 and ICAM1 molecules for 20 min. Scale bar, 10 μM (**c**) Human T_sen_ have defects in FAO, which are reversed upon CPT1A rescue. Senescent T cells were activated with anti-CD3 and recombinant human IL-2 for 48 hours followed by transduction with mock (CTRL) or CPT1A lentiviral vector (*n* = 3 donors). FAO rate was assessed by ELISA assays in transduced T cells 96 hours later. Data are expressed as fold change to non-senescent T_erl_, set as 1. (**d**) T_sen_ generate ceramides from palmitate in the absence of FAO. ELISA-based *de novo* ceramide synthesis in T_sen_ cells transduced with CTRL or CPT1A vectors, treated with or without SPT inhibitor myriocin (10 μM) then stimulated overnight in the presence or absence of palmitate (100 μM). Ceramide levels were assessed by anti-ceramide antibody coupled to ELISA (*n* = 3 donors). Data are expressed as fold change to CTRL cells, set as 1. (**e**) Ceramidase activity assessed by ELISA-based assays. Human ceramidase was immunoprecipitated with anti-ASAH2 antibody directly *ex vivo* from T_sen_ vs T_erl_ (*n* = 3 donors; **right**) or from T_sen_ cells transduced as indicated and then incubated (1 hour at 37 °C) with different doses [50 μM or 2000 μM] of C-12 ceramides (*n* = 3 donors; **left**) followed by monitoring of ceramide conversion into sphingosine. The conversion reaction into sphingosine was monitored by fluorescence emission at 450 nm after addition of o-phthaldialdehyde (o-PA). Note that increased concentration of ceramides inhibits ceramidase activity. Data are expressed as fold change to CTRL or T_erl_ cells, set as 1. CE, ceramide. (**f**) Representative flow-cytometry showing that T_sen_ express lower phosphatidylethanolamine (PE) as compared to non-senescent T_erl_ on their membrane. PE levels were assessed by PE-specific duramycin staining (*n* = 3 donors). (**g**) Synapse-formation on planar bilayers showing that CPT1A-driven FAO restoration requires ceramidase activation and subsequent phosphatidylethanolamine production. As such, T_sen_ transduced with CPT1A-expressing vectors were treated with ceramidase inhibitors for 10 days, rested overnight, then treated with or without phosphatidylethanolamine (PE; 2 μM 20 min) prior to 20 min stimulation on bilayers. Representative of *n* = 4 donors. Scale bars, 10 μM. Pooled data (*n* = 37 synapses per condition) (**right**). Paired student’s t test in (**b, e right, g**), ANOVA with Bonferroni-post test correction in (**c, d, e left**). *P < 0.05, **P < 0.01, ***P < 0.001, ****P < 0.0001. Error bars indicate s.e.m.

**Extended Data Fig. 2:**
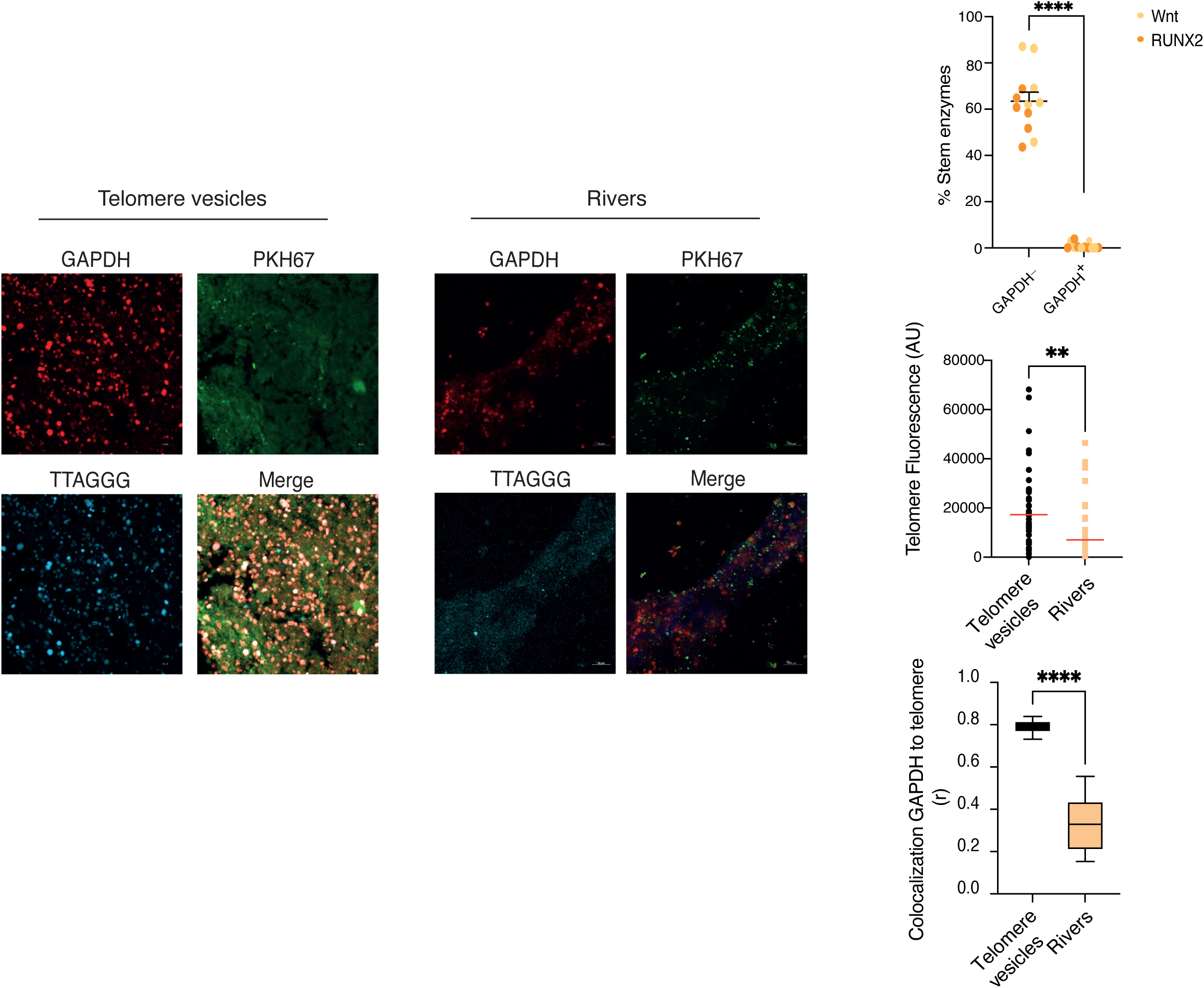
Relationship of APC GAPDH with telomere Rivers. Flow cytometry and Immunofluorescence with telomere FISH coupled to anti-GAPDH detection. Telomere vesicles (left) or River particles (right) were subjected to differential centrifugation and analyzed as indicated. Individual telomeres are shorter in the Rivers yet the particles are bigger in size likely entrapped in NET-like structures^28^. Note that GAPDH depleted telomere Rivers possess stemness factors (*n* = 12 donors; top, right). Fluorescence of telomere vesicles *vs* River-like particles assessed by IF-FISH (middle, right). Pearson’s co-localization (r) of GAPDH to telomeres is shown (bottom, right), demonstrating that telomere Rivers are largely devoid of GAPDH, in contrast to the telomere vesicles. Data are from 9 microscopy images (*n* = 3 donors). Scale bars, 10 μM. Mann-Whitney test was used for statistical analysis. **P < 0.01, ****P < 0.0001. Error bars indicate s.e.m.

**Extended Data Fig. 3:**
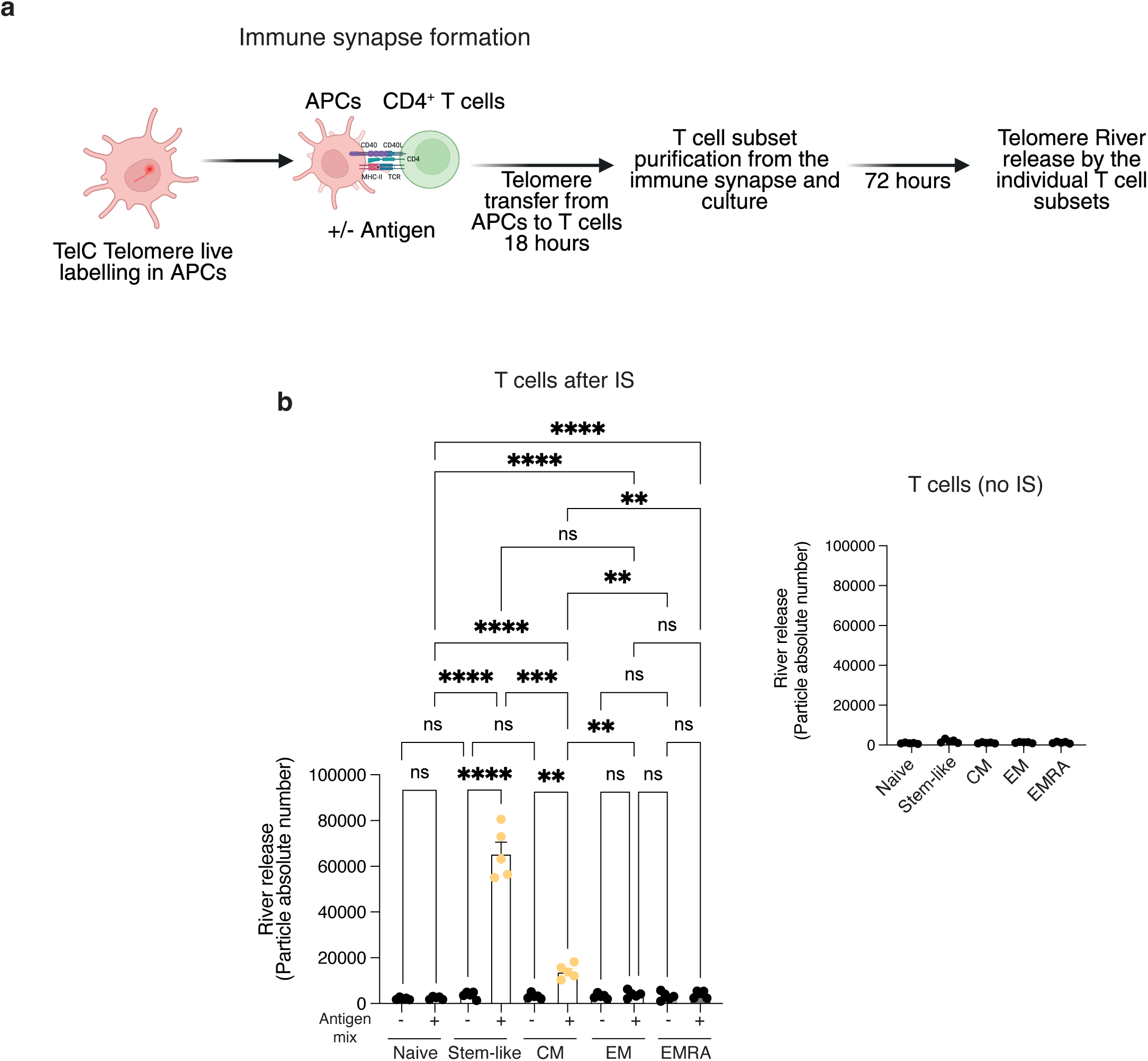
Human CD4⁺ T cell subsets differ in River production capacity. (**a**) Experimental design. Total human CD4⁺ T cells were conjugated with autologous APCs pre-labelled with TelC and pulsed with or without an antigen mix (influenza, CMV, EBV). At 24 h post-conjugation, CD4⁺ T cells were FACS-sorted into naïve (CD27⁺CD28⁺CD45RA⁺), central memory (CD27⁺CD28⁺CD45RA⁻), effector memory (CD27⁻CD28⁻CD45RA⁻), and TEMRA (CD27⁻CD28⁻CD45RA⁺) subsets, then cultured an additional 48 h without further stimulation. Supernatants were collected for River detection. (**b**) River detection in culture supernatants. Naïve and central memory produced Rivers (PKH67⁺TelC⁺GAPDH⁻), whereas effector memory/TEMRA did not. Antigen-free APCs gave background only. Sorted T cells without APCs released no Rivers, confirming antigen dependency. Pooled data (*n* = 6 donors) are shown. In control experiments, T-cell subsets sorted and cultured without APCs or activation released no Rivers, demonstrating T-cell activation as essential for River production. One-way ANOVA with Bonferroni post-test. ***p < 0.001, ****p < 0.0001. Error bars, s.e.m.

**Extended Data Fig. 4:**
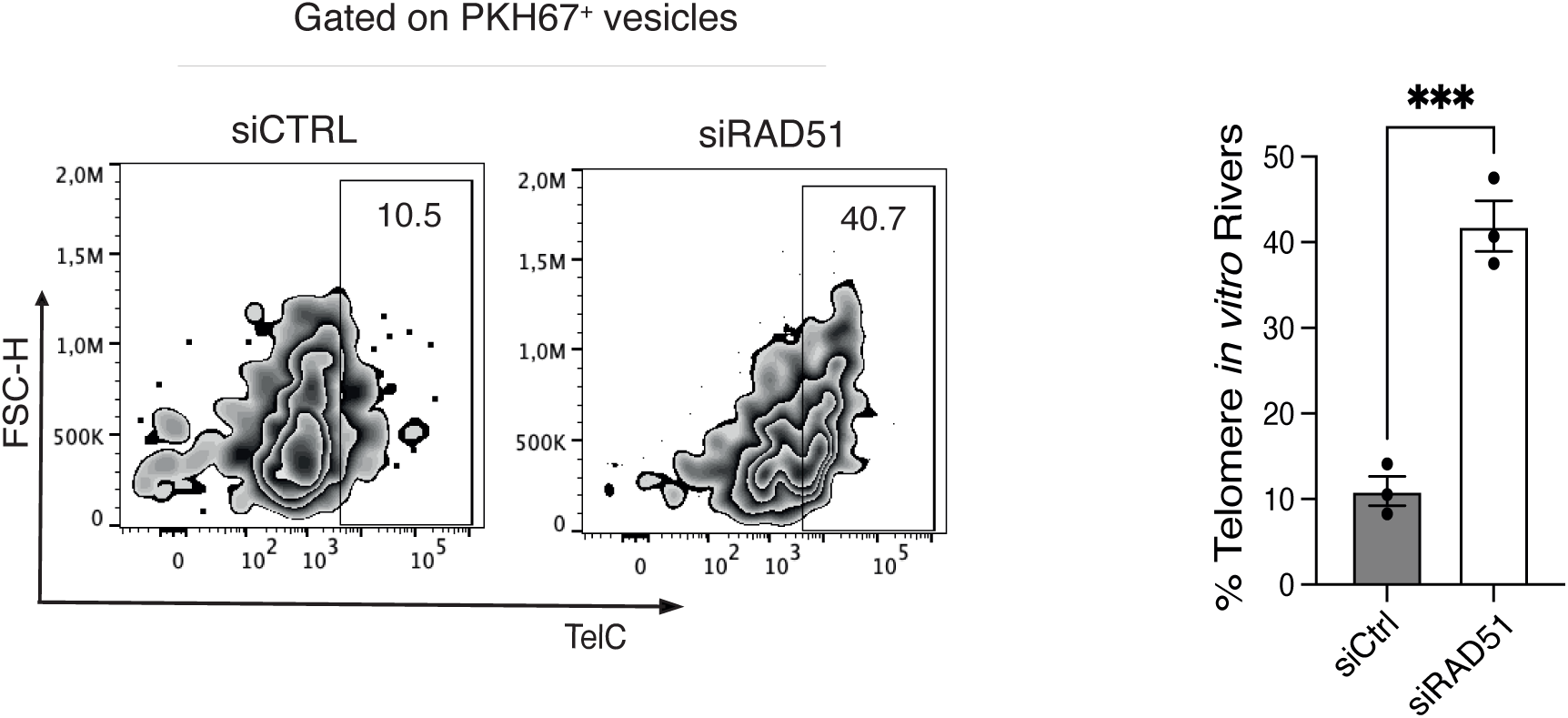
Telomere Rivers are composed of supernumerary APC telomeres. siCTRL or siRAD51 telomere vesicles were derived by ionomycin APC activation and provided to T cells, followed by anti-CD3 and anti-CD28 activation for 48 hours. T cell supernatants were then assessed for presence of River-like particles. Note that suppression of APC transfer-driven T cell telomere elongation augmented River release (*n* = 3 donors). Paired student’s t test was used. ***P < 0.001. Error bars indicate s.e.m.

**Extended Data Fig. 5:**
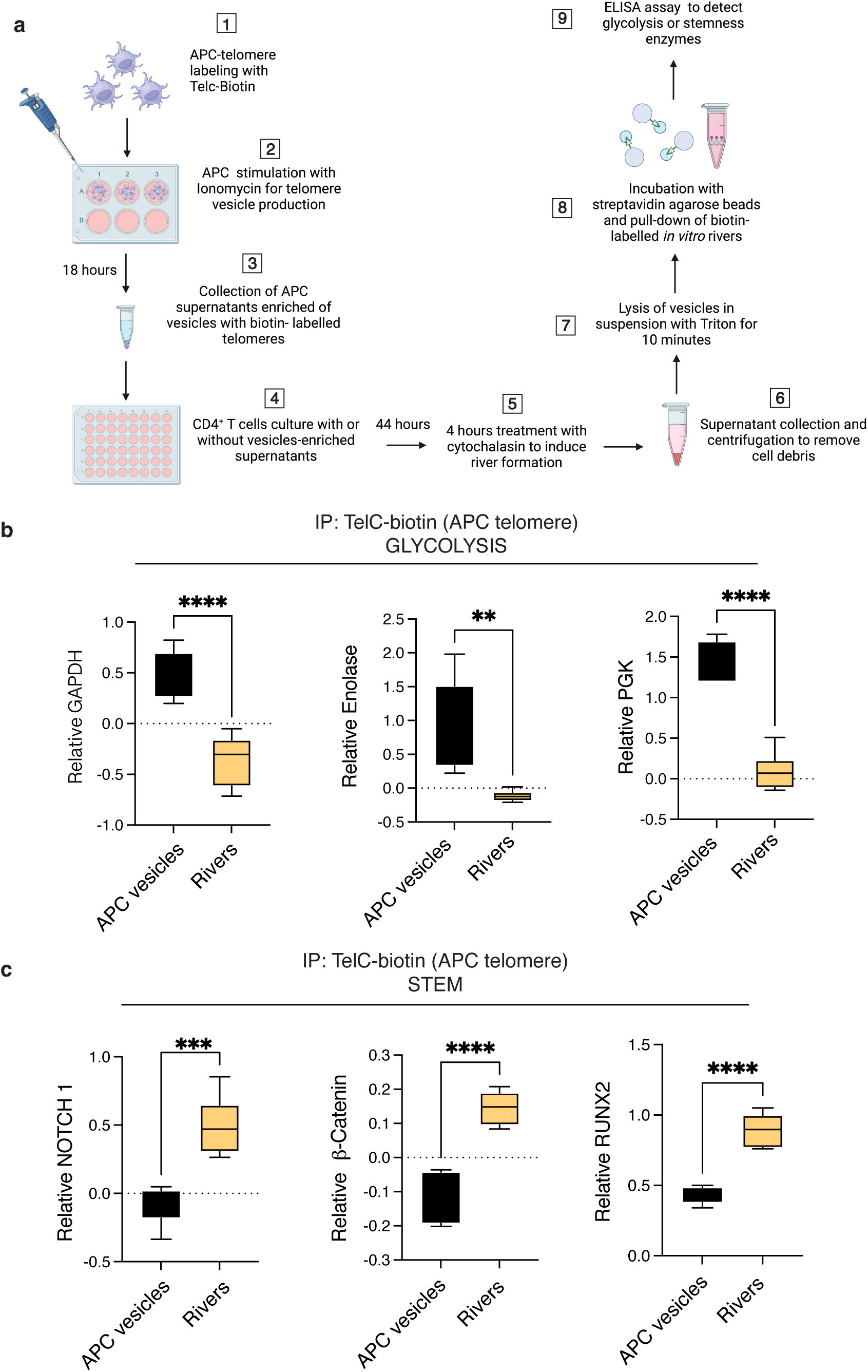
Composition of telomere Rivers, biochemical evidence. (**a**) Experimental design. Human APCs were pre-labelled with TelC-biotin probes, stimulated with ionomycin, and their telomeres subsequently transferred into autologous CD4⁺ T cells. Co-cultures were maintained for 48 h, with cytochalasin D added 4 h before termination. Culture supernatants were collected and subjected to streptavidin agarose immunoprecipitation, and precipitated APC-derived telomeres were analyzed by ELISA. APC telomere vesicles were analyzed in parallel as controls. (**b**) Presence of glycolytic enzymes (GAPDH, Enolase and PGK) in APC telomere vesicles or River-like particles derived from the purified T cell cultures and (**c**) existence of stem enzymes (NOTCH1, β-Catenin and RUNX2) in the same conditions. Data are from *n* = 6 donors. Unpaired student’s t test in (**b**, **c**). **P < 0.01, ***P < 0.001, ****P < 0.0001. Error bars indicate s.e.m.

**Extended Data Fig. 6:**
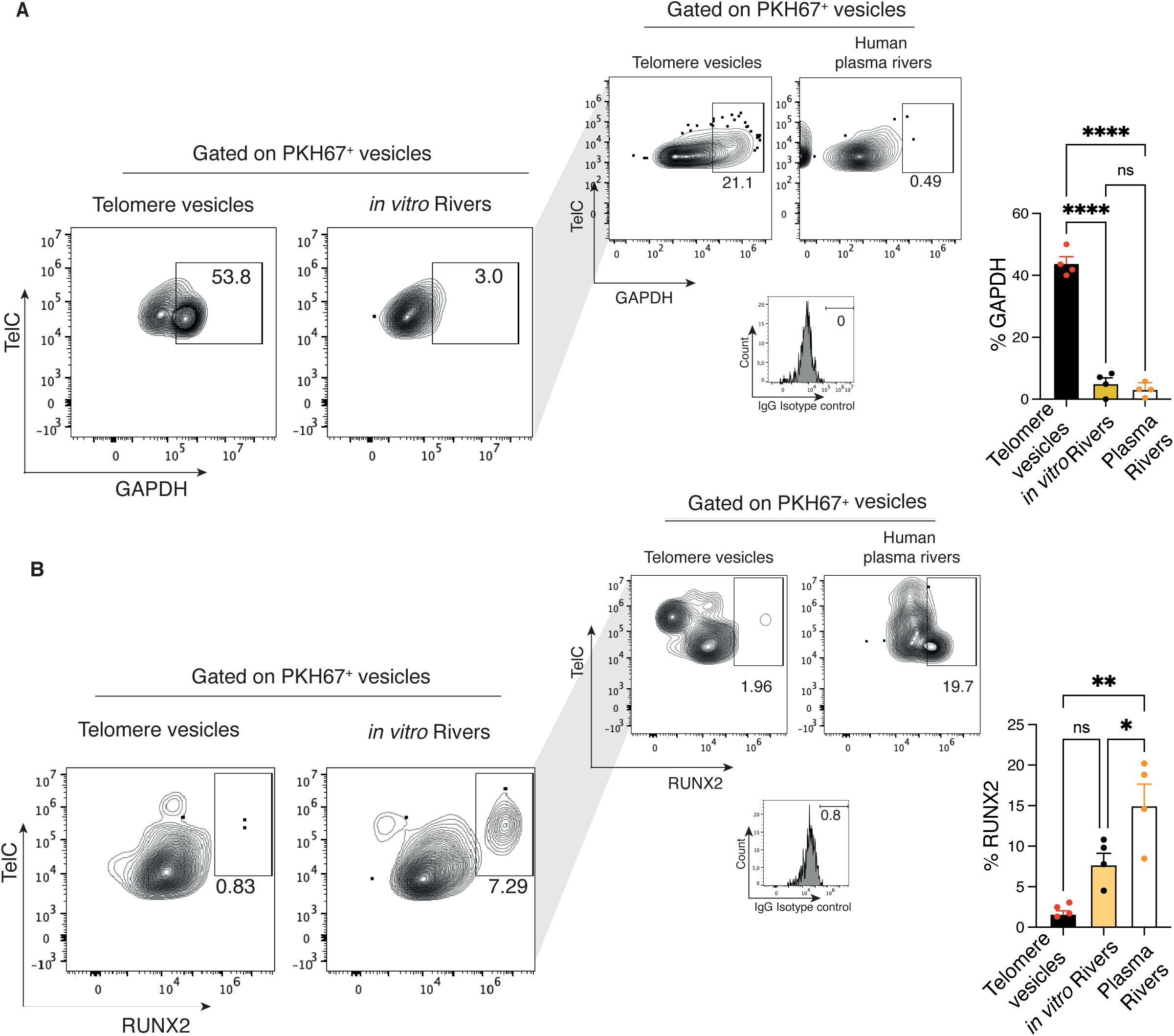
Comparison of telomere Rivers *in vitro* and in human sera. Flow-cytometry based assessment of (**a**) GAPDH among telomere vesicles or *in vitro* telomere Rivers as per Extended data 4 or in human sera. River composition is defined as PKH67^+^ TelC^+^ and GAPDH^low/-^particles. The same assessment for stem-related RUNX2 enzyme is shown (**b**). Isotype controls are shown. Note the similarities between *in vitro* generated Rivers and those naturally circulating in humans. Data are from *n* = 4 donors. One-way Anova with Bonferroni post-correction for multiple comparisons. *P < 0.05, **P < 0.01, ****P < 0.0001. Error bars indicate s.e.m.

**Extended Data Fig. 7:**
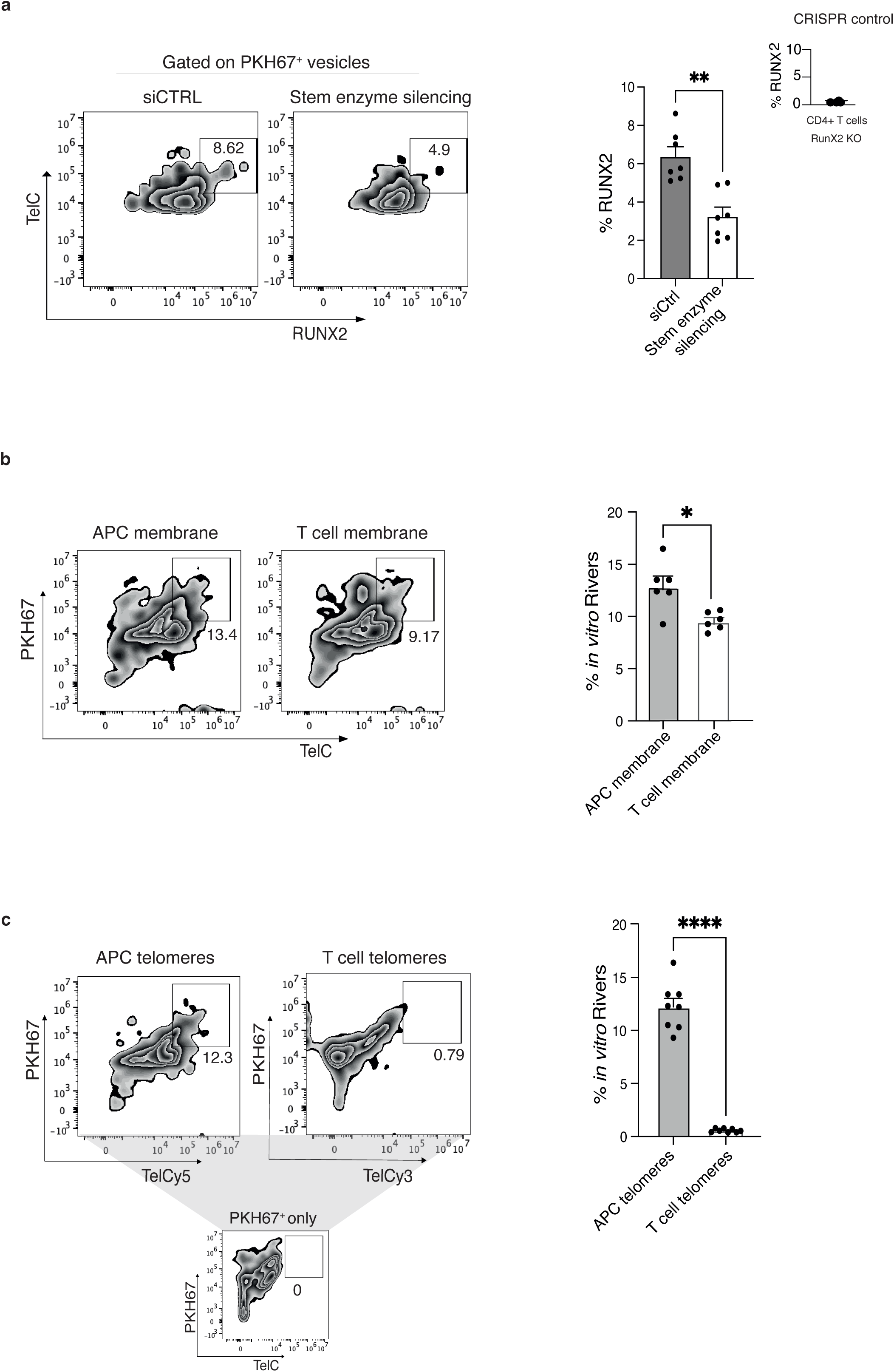
River composition controls. (**a**) T cell origin of stem-associated enzymes. CD4⁺ T cells were transfected with siCTRL or a triple stem-enzyme knockdown (siNOTCH1, siWnt5, siRUNX2). Cells were then co-cultured with fluorescently labelled APC telomere vesicles and activated with anti-CD3/CD28 for 48 h. Supernatants were analyzed for the presence of stem enzymes in River-like particles. Knockdown of T cell enzymes abolished stem-related protein content in Rivers without affecting absolute particle numbers. Data are from (*n* = 7 donors). (**b**) APC or T cell contribution to River membranes. Representative FACS plots (left) and pooled quantification (right) of Rivers generated *in vitro* from APCs or T cells pre-labelled with PKH67 membrane dye. APC telomere vesicles were derived by ionomycin stimulation and transferred to T cells, followed by 48 h activation with anti-CD3/CD28. Both APC- and T cell–derived lipids were incorporated into River membranes (*n* = 6 donors). (**c**) APC or T cell contribution to River telomeres. Representative FACS plots (left) and quantification (right) of Rivers generated in vitro from APCs or CD4⁺ T cells pre-labelled with TelC-Cy5 or TelC-Cy3 probes, respectively. Rivers were gated as PKH67⁺TelC⁺. APC telomeres (TelC-Cy5) were consistently detected in Rivers, whereas no T cell telomeres (TelC-Cy3) were incorporated (*n* = 8 donors). TelC-free controls are shown. Paired student’s t test in (**a**-**c**). *P < 0.05, **P < 0.01, ****P < 0.0001. Error bars indicate s.e.m.

**Extended Data Fig. 8:**
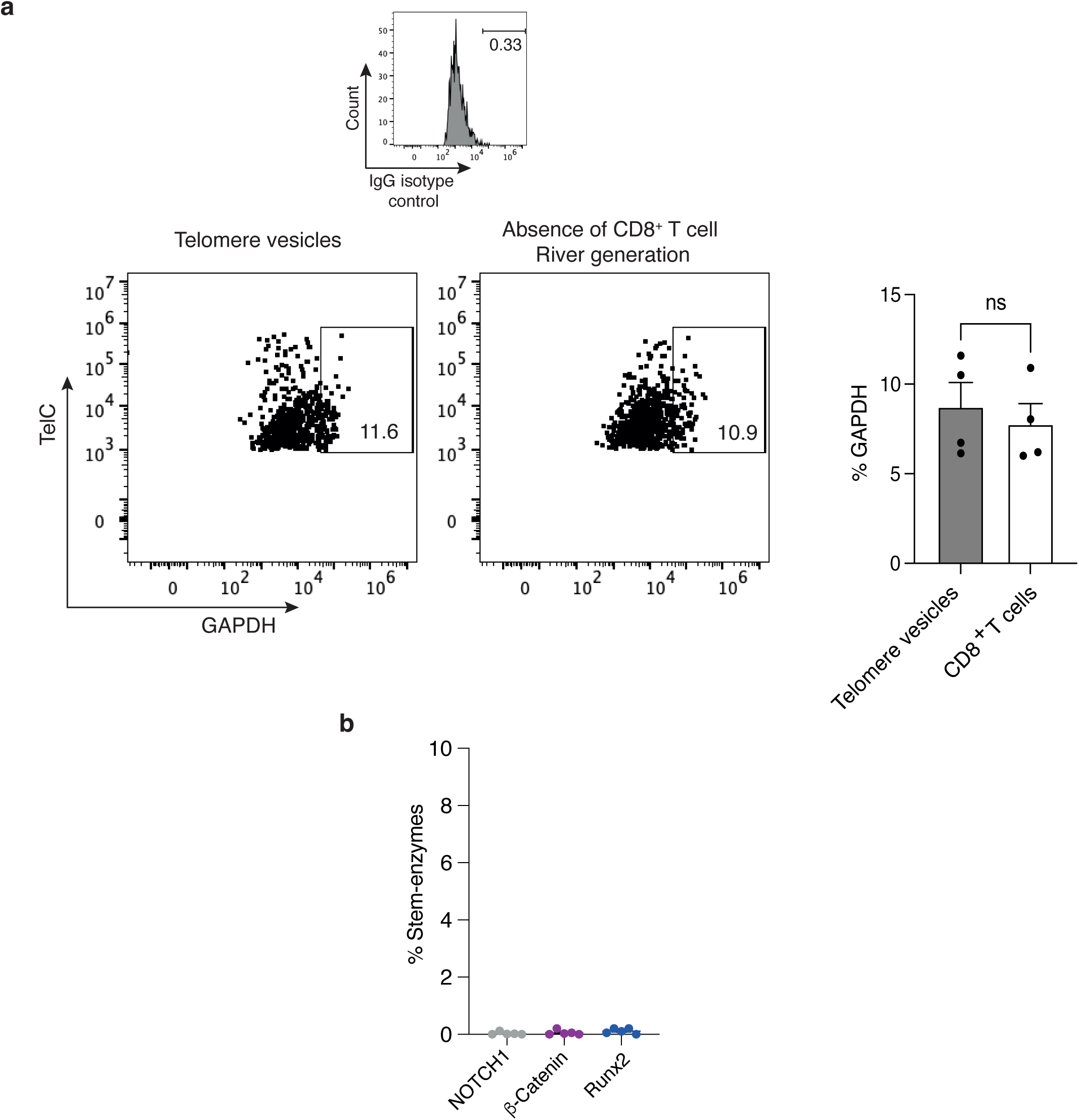
CD8^+^ T cells do not generate Rivers. Human CD8⁺ T cells were incubated with fluorescently labelled APC telomere vesicles and activated with anti-CD3/CD28 for 48 h. Culture supernatants were analyzed for GAPDH clearance from telomere vesicles as evidence of River-like particle generation. In parallel, stem-associated markers (NOTCH1, β-Catenin, RUNX2) were assessed to test for stem enrichment. Unlike CD4⁺ T cells, CD8⁺ T cells failed to clear GAPDH or enrich stem proteins, indicating inability to generate *bona fide* Rivers. Representative FACS plots and isotype controls are shown. Data from (*n* = 4 donors). Statistical analysis by paired Student’s t test; ns, non-significant.

**Extended Data Fig. 9:**
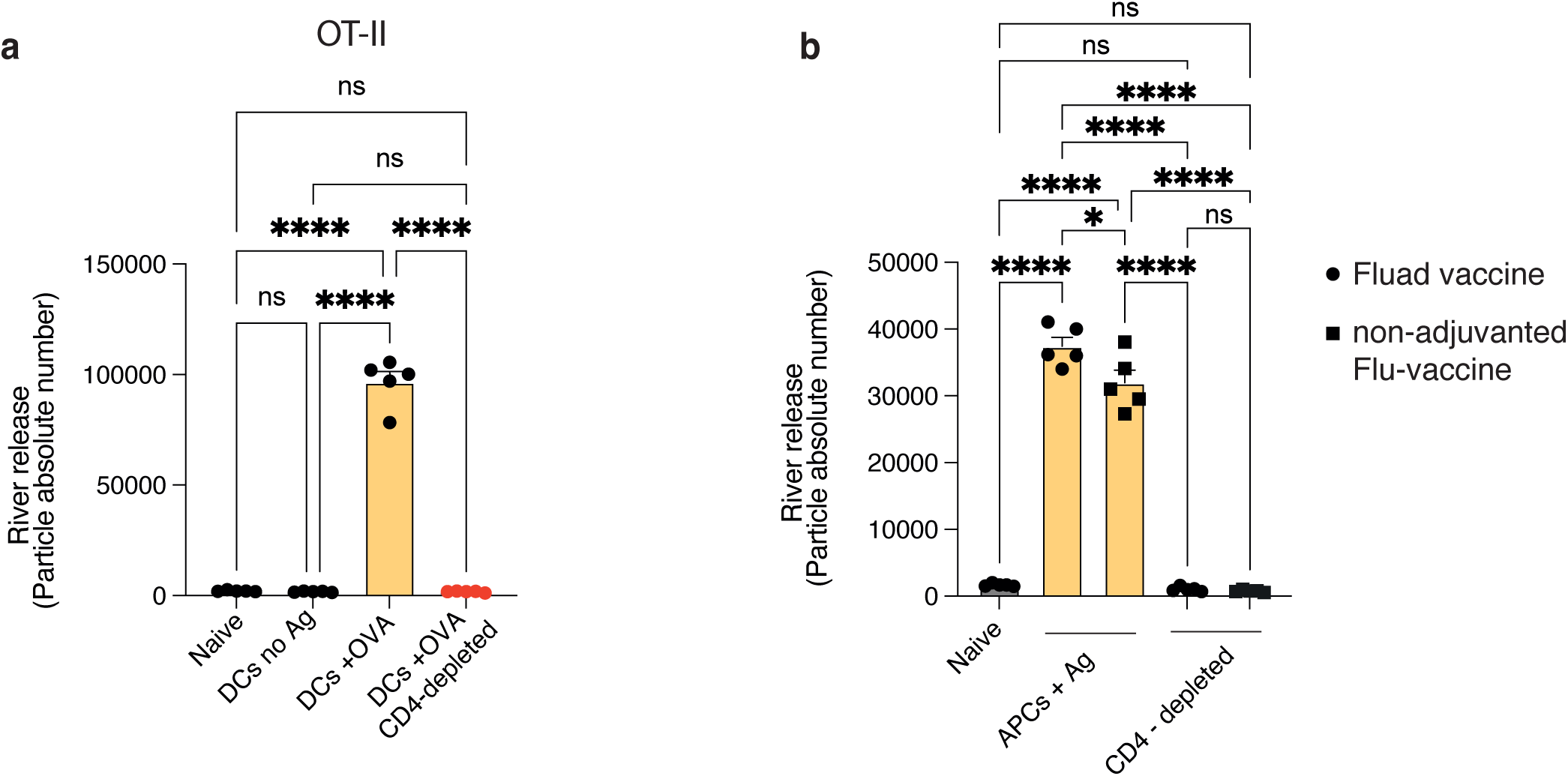
Antigen-driven River generation is independent of adjuvant. (**a**) Splenic CD11c⁺ dendritic cells were purified from young C57BL/6 mice, labelled with TelC, and pulsed with OVA peptide (OT-II 323–339). Cells were transferred subcutaneously into OT-II transgenic recipients with or without prior CD4⁺ T cell depletion. Serum was collected at day 15 and Rivers were quantified as PKH67⁺TelC⁺GAPDH⁻ by flow cytometry/FAVS. Robust River formation occurred after transfer of OVA-pulsed DCs in CD4-proficient recipients, whereas CD4 depletion abolished River production. (**b**) Splenic APCs (CD3-) were labelled with TelC and pulsed with influenza vaccine, either adjuvanted (Fluad) or non-adjuvanted (Fluarix), then transferred subcutaneously into young C57BL/6 mice. In parallel, some hosts were depleted of CD4⁺ T cells. Serum was collected at day 15 and Rivers were quantified as PKH67⁺TelC⁺GAPDH⁻ by flow cytometry/FAVS. Data are from (*n* = 5 mice per group). One-way ANOVA with Bonferroni correction. **\*\*\*\***P < 0.0001. Error bars indicate s.e.m.

**Extended Data Fig. 10:**
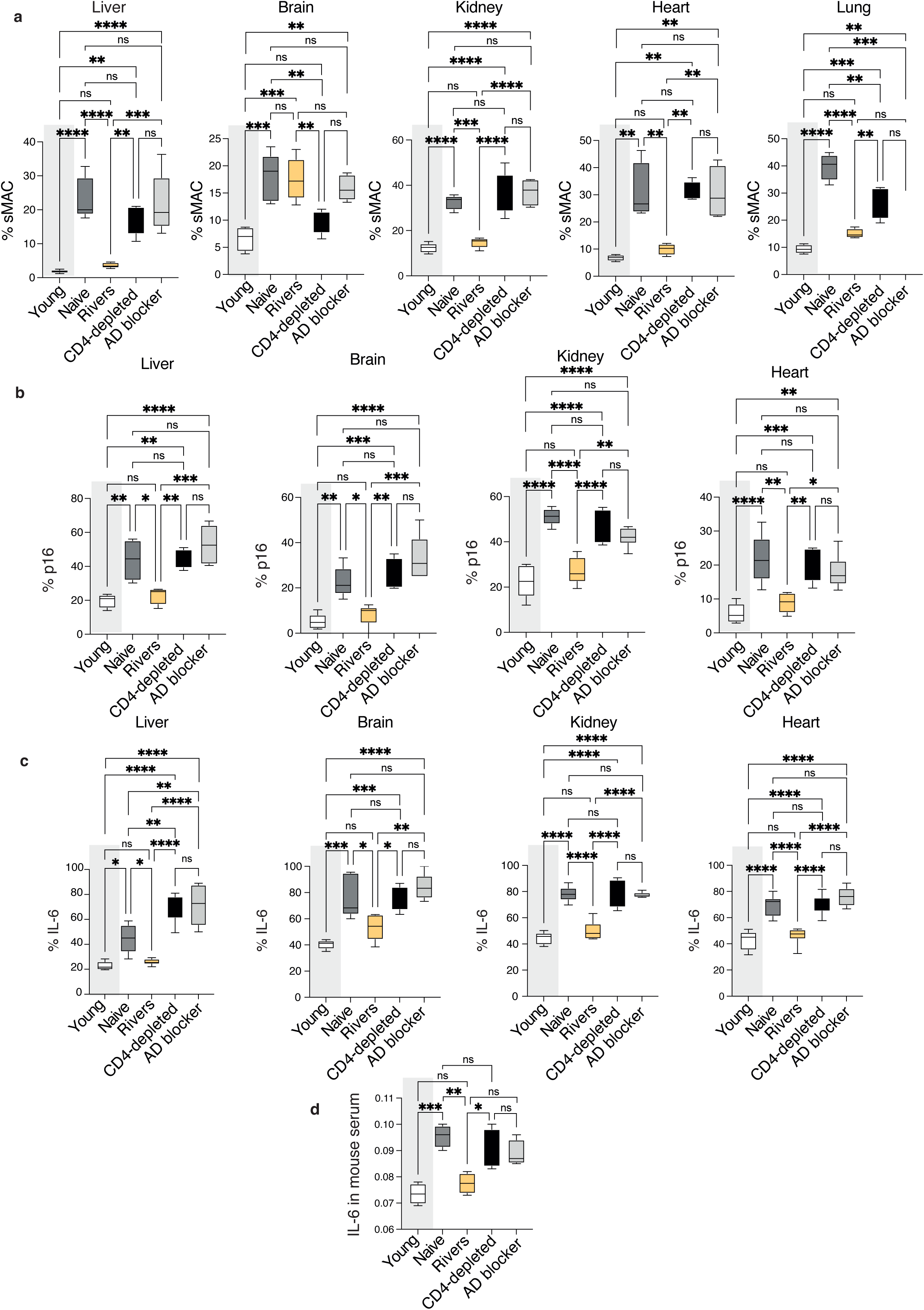
Systemic rejuvenation by River transplant. Old recipient mice received subcutaneous River injections then bred normally for 12 months. Peripheral organs (Liver, Brain, Kidney, Heart and Lung) were assessed for senescence-associated markers (sMAC (**a**), p16 (**b**), IL-6 (**c**)) by flow-cytometry. Note that only Rivers generated in the absence of anti-LFA1 antibody or CD4^+^ T cell depletion are effectively produced (Fig. 4) and as such rejuvenate tissues. Young (3-month-old) mice are shown, as control. (**d**) Assessment of the SASP factor IL-6 in sera of animals as in (a-c). Note that River transplant abolishes senescence and inflammatory markers. In **a**, data are from *n* = 4 mice (hearth, lung) or *n* = 5 mice (liver, brain, kidney) per group; in **b**, data are from *n* = 5 mice (liver, brain) or *n* = 6 mice (kidney, hearth) per group; in **c**, data are from *n* = 5 mice (brain) or *n* = 6 mice (liver, kidney), or *n* = 7 mice (hearth) per group; in **d**, data are from *n* = 4 mice per group. In (**a**-**d**), One-way Anova with Bonferroni post-correction for multiple comparisons. *P < 0.05, **P < 0.01, ***P < 0.001, ****P < 0.0001. Error bars indicate s.e.m.

**Extended Data Fig. 11:**
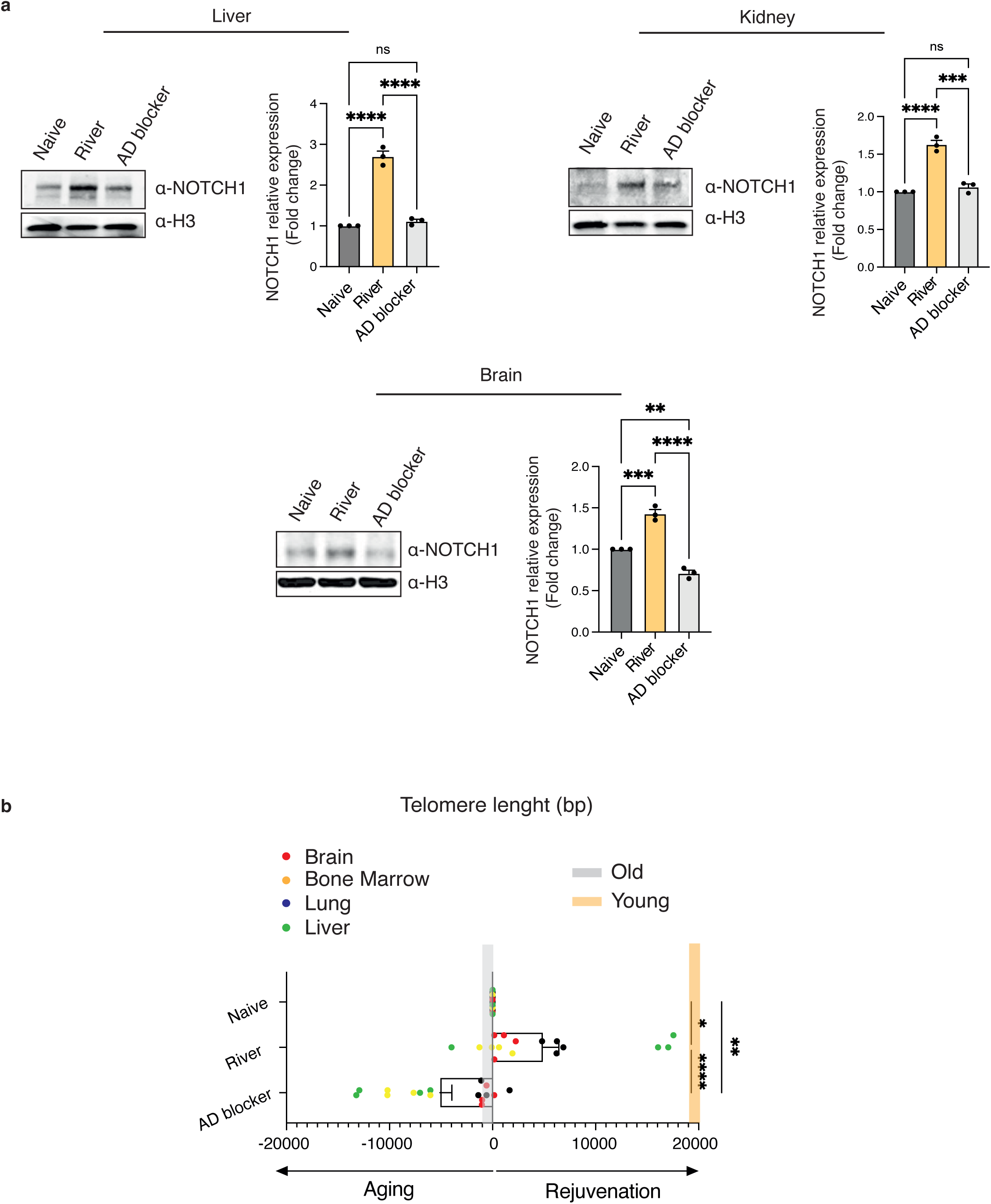
Stem induction and telomere extension in River irrigated tissues. (**a**) Immunoblots of stem-related NOTCH1 in liver, kidney and brain of mice one year after River transplant as per Fig. 4b, c. Representative immunoblots and pooled data from *n* = 3 mice per group are shown. (**b**) Flow-FISH of mice as in **a** for the organs shown from 3 mice per group (*n* = 12 organs). In (**a**, **b**) One-way Anova with Bonferroni post-correction for multiple comparisons. *P < 0.05, **P < 0.01, ***P < 0.001, ****P < 0.0001. Error bars indicate s.e.m.

**Extended Data Fig. 12:**
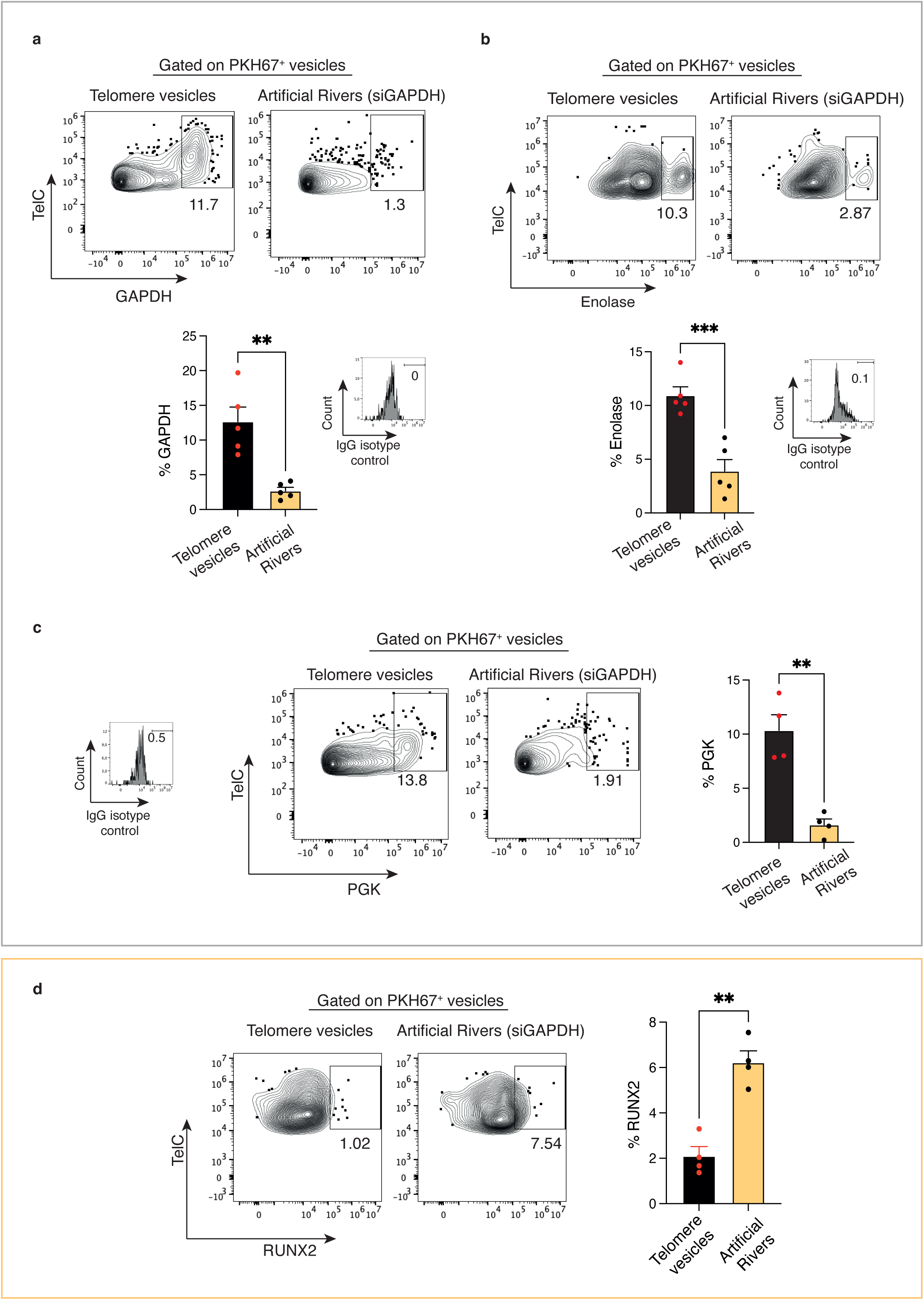
Artificial River characterization. Absence of glycolysis enzymes in artificial Rivers, phenocopying natural Rivers. FACS plots and pooled data showing that silencing of APC GAPDH (**a**) generates artificial Rivers that lack enolase (**b**) and Phosphoglycerate kinase (PGK) (**c**) glycolysis enzymes. Data are obtained by gating on PKH67^+^ and TelC^+^ APC vesicles and are from *n* = 5 mice (**a -b**) or *n* = 4 mice (**c-d**) per group. IgG isotype controls are shown. Presence of stem enzymes (RUNX2) in artificial Rivers derived and analyzed as in **a**. In (**a**-**d**) paired student’s t test. **P < 0.01, ***P < 0.001. Error bars indicate s.e.m.

## References

1. Conese, M., Carbone, A., Beccia, E. & Angiolillo, A. The Fountain of Youth: A tale of parabiosis, stem cells, and rejuvenation. Open Medicine 12, 376–383 (2017).

2. Pálovics, R. et al. Molecular hallmarks of heterochronic parabiosis at single-cell resolution. Nature 603, 309–314 (2022).

3. Reardon, S. ‘Young blood’ anti-ageing mechanism called into question. Nature (2015).

4. Diggs, J. Cellular Theory of Aging. Encyclopedia of Aging and Public Health, 1981–99 (2008).

5. Lanna, A. et al. A sestrin-dependent Erk-Jnk-p38 MAPK activation complex inhibits immunity during aging. Nat Immunol 18, 354–363 (2017).

6. Lanna A et al. An intercellular transfer of telomeres rescues T cells from senescence and promotes long-term immunological memory. Nat Cell Biol 24, 1461–1474 (2022).

7. Pearce, E. L. et al. Enhancing CD8 T-cell memory by modulating fatty acid metabolism. Nature 460, 103–7 (2009).

8. van der Windt, G. J. W. et al. Mitochondrial Respiratory Capacity Is a Critical Regulator of CD8+ T Cell Memory Development. Immunity 36, 68–78 (2012).

9. Kunz, H.-H. et al. The ABC Transporter PXA1 and Peroxisomal b-Oxidation Are Vital for Metabolism in Mature Leaves of Arabidopsis during Extended Darkness. Plant Cell 21, 2733–2749 (2009).

10. Chen, L. et al. Fenofibrate-Induced Mitochondrial Dysfunction and Metabolic Reprogramming Reversal: The Anti-Tumor Effects in Gastric Carcinoma Cells Mediated by the PPAR Pathway. Am J Transl Res 12 (2020).

11. Hasanpourghadi, M. et al. Treatment with the PPARα agonist fenofibrate improves the efficacy of CD8+ T cell therapy for melanoma. Mol Ther Oncolytics 31, (2023).

12. Desdín-Micó, G. et al. T cells with dysfunctional mitochondria induce multimorbidity and premature senescence. Science 368, 1371–1376 (2020).

13. Henson, S. M. et al. P38 signaling inhibits mTORC1-independent autophagy in senescent human CD8+ T cells. Journal of Clinical Investigation 124, 4004–4016 (2014).

14. Perry, D. K. et al. Serine Palmitoyltransferase Regulates *de novo* Ceramide Generation during Etoposide-induced Apoptosis. Journal of Biological Chemistry 275, 9078–9084 (2000).

15. Varma, R., Campi, G., Yokosuka, T., Saito, T. & Dustin, M. L. T Cell Receptor-Proximal Signals Are Sustained in Peripheral Microclusters and Terminated in the Central Supramolecular Activation Cluster. Immunity 25, 117–127 (2006).

16. Choudhuri, K. et al. Polarized release of T-cell-receptor-enriched microvesicles at the immunological synapse. Nature 507, 118–23 (2014).

17. MacEyka, M. & Spiegel, S. Sphingolipid metabolites in inflammatory disease. Nature 510, 58–67 (2014).

18. Calzada, E., Onguka, O. & Claypool, S. M. Phosphatidylethanolamine Metabolism in Health and Disease. Int Rev Cell Mol Biol 321, 29–88 (2016).

19. Vaena, S. et al. Aging-dependent mitochondrial dysfunction mediated by ceramide signaling inhibits antitumor T cell response. Cell Rep 35, 109076 (2021).

20. Doerr, A. DIA mass spectrometry. Nature Methods 12, 35 (2014).

21. Lazarev, V. F., Guzhova, I. V. & Margulis, B. A. Glyceraldehyde-3-phosphate dehydrogenase is a multifaceted therapeutic target. Pharmaceutics 12 (2020).

22. Sunchu, B. & Cabernard, C. Principles and mechanisms of asymmetric cell division. Development 147 (2020).

23. Chang J T et al. Asymmetric T lymphocyte division in the initiation of adaptive immune responses. Science 315, 1687–1691 (2007).

24. Verbist, K. C. et al. Metabolic maintenance of cell asymmetry following division in activated T lymphocytes. Nature 532, 389–393 (2016).

25. Ito, K. et al. A PML-PPAR-δ pathway for fatty acid oxidation regulates hematopoietic stem cell maintenance. Nat Med 18, 1350–1358 (2012).

26. Pollizzi, K. N. et al. Asymmetric inheritance of mTORC1 kinase activity during division dictates CD8+ T cell differentiation. Nat Immunol 17, 704–711 (2016).

27. Catalano, M. & O’Driscoll, L. Inhibiting extracellular vesicles formation and release: a review of EV inhibitors. Journal of Extracellular Vesicles 9 (2020).

28. Ebrahimi, F. et al. Markers of neutrophil extracellular traps predict adverse outcome in communityacquired pneumonia: Secondary analysis of a randomised controlled trial. European Respiratory Journal 51, 1701389 (2018).

29. Moriyama, H. et al. Notch signaling enhances stemness by regulating metabolic pathways through modifying p53, nf-κb, and hif-1α. Stem Cells Dev 27, 935–947 (2018).

30. Valenti, M. T. et al. Runx2 expression: A mesenchymal stem marker for cancer. Oncol Lett 12, 4167–4172 (2016).

31. Gattinoni, L. et al. Wnt signaling arrests effector T cell differentiation and generates CD8 + memory stem cells. Nat Med 15, 808–813 (2009).

32. Capece, T. et al. A novel intracellular pool of LFA-1 is critical for asymmetric CD8+ T cell activation and differentiation. Journal of Cell Biology 216, 3817–3829 (2017).

33. Dimri, G. P. et al. A biomarker that identifies senescent human cells in culture and in aging skin in vivo. Proc Natl Acad Sci U S A 92, 9363–9367 (1995).

34. Gorgoulis, V. et al. Cellular Senescence: Defining a Path Forward. Cell 179, 813–827 (2019).

35. Sun, Y., Li, Q. & Kirkland, J. L. Targeting senescent cells for a healthier longevity: the roadmap for an era of global aging. Life Medicine 1, 103–119 (2022).

36. Lanna, A., Henson, S. M., Escors, D. & Akbar, A. N. The kinase p38 activated by the metabolic regulator AMPK and scaffold TAB1 drives the senescence of human T cells. Nat Immunol 15, 965–72 (2014).

37. Lanna, A. et al. Rejuvenation driven reprograming in T lymphocytes. *In Review*, Nature. doi:10.21203/rs.3.rs-5180379/v1

38. Polonsky M., et al. Induction of CD4 T cell memory by local cellular collectivity. Science 360, eaaj1853 (2018).

39. Amor, et al., Prophylactic and long-lasting efficacy of senolytic CAR T cells against age-related metabolic dysfunction. Nat Aging 4, 336–349 (2024).

40. Liu M N.., Lan Q., Wu H., Qiu C.W. Rejuvenation of young blood on aging organs: Effects, circulating factors, and mechanisms. Heliyon, 10 e32652 (2024).

41. Jaskelioff, M., et al. Telomerase reactivation reverses tissue degeneration in aged telomerase-deficient mice. Nature 469, 102–106 (2011).

