## Supplementary information; Lanna et al 2025 for "CD4⁺ T cells confer transplantable rejuvenation via Rivers of telomeres"

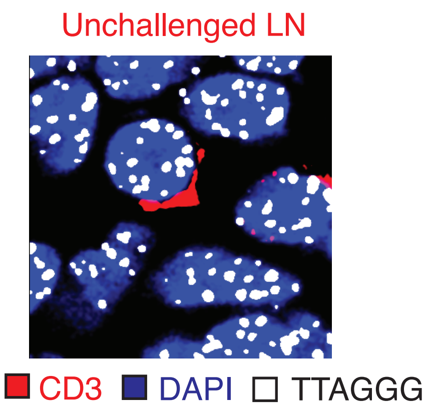


**Supplementary information 1:** Absence of extracellular telomeres in lymph node vessels from naive (young) animals not subjected to vaccination. A representative T cell with intracellular telomeres is shown.

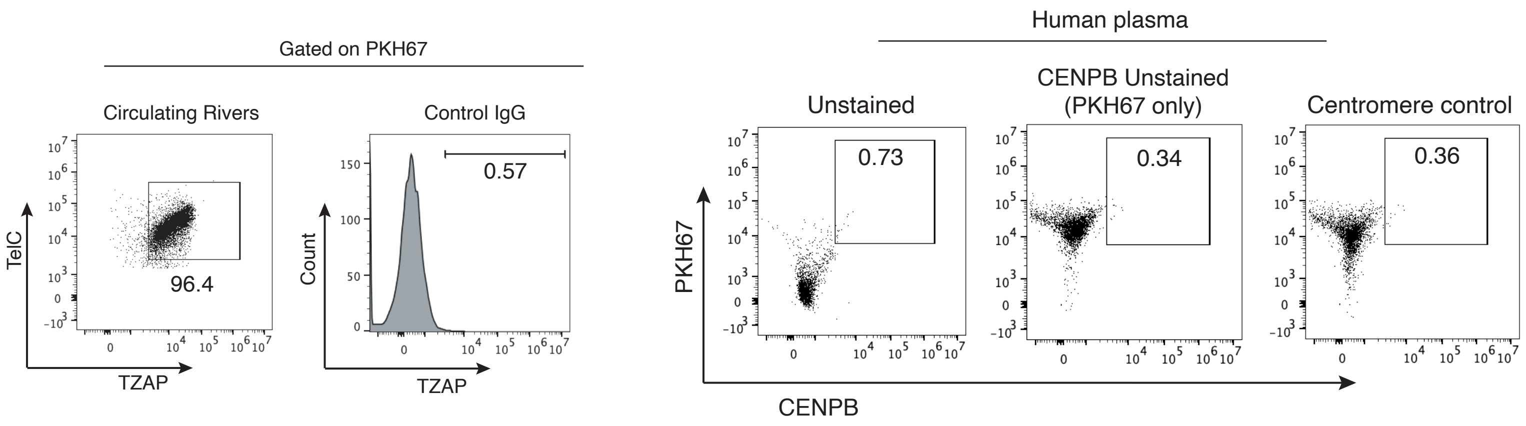


**Supplementary information 2:** TZAP staining of PKH67⁺ TelC⁺ human circulating Rivers. Note that all TelC⁺ River particles express the telomere-binding factor TZAP⁺ (left). Representative FACS plot showing human circulating RIvers stained with PKH67 dye and with centromere protein B probe (CENPB). Unstained and PKH67 single-stained controls were used to define gating thresholds (right).

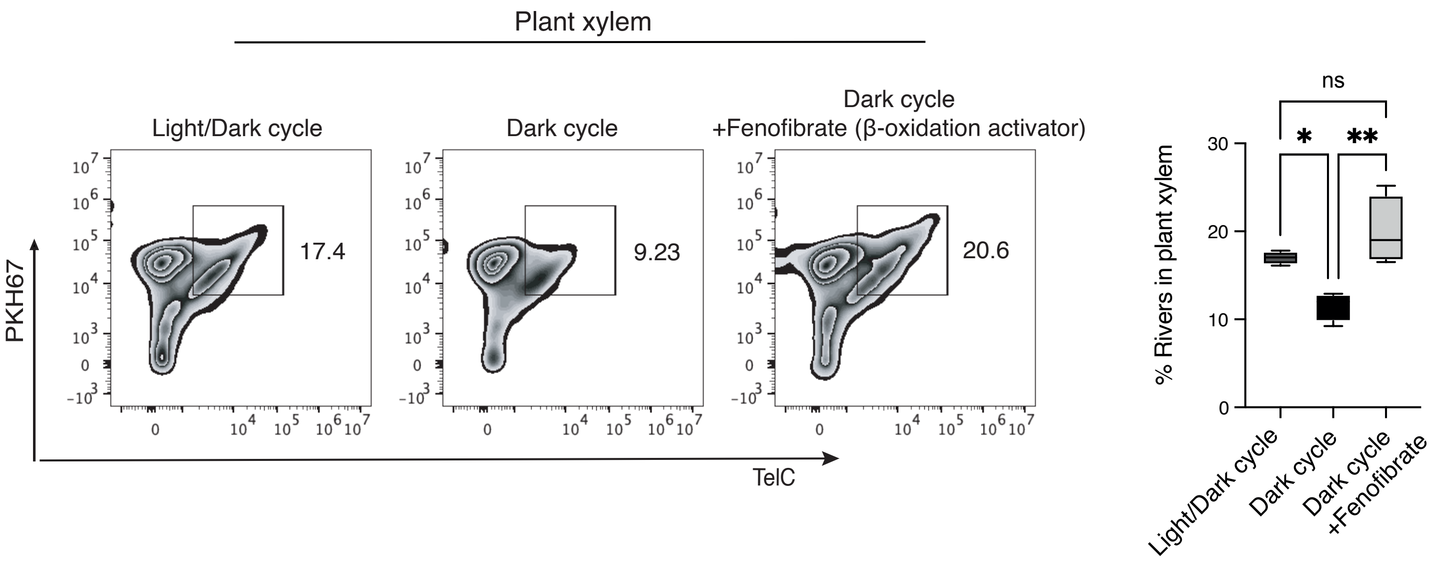


**Supplementary information 3:** Hydration of plant roots with fenofibrate restores telomere River production in plant xylem, lost when a dark/dark cycle is enforced.

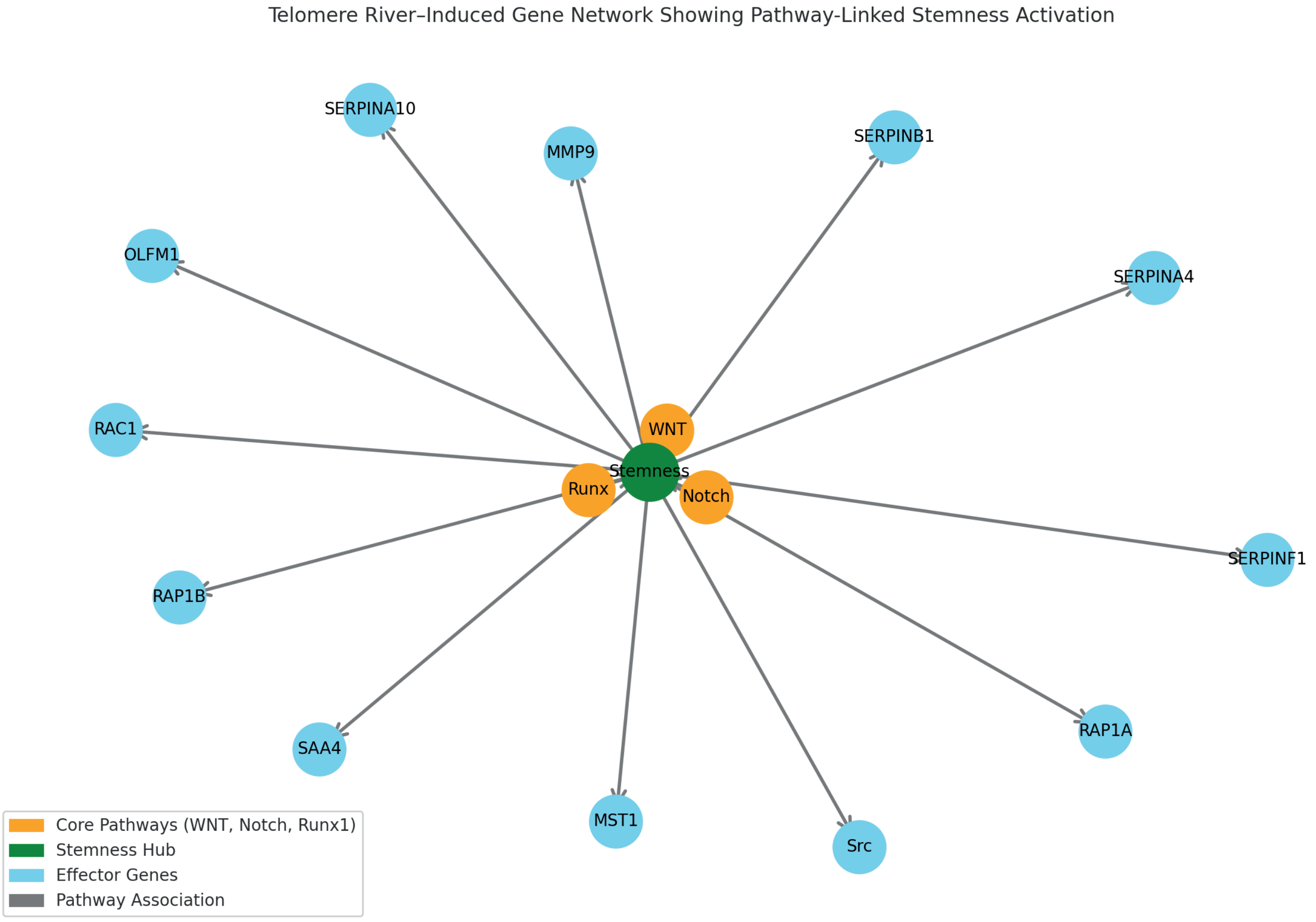


**Supplementary information 4.** Gene set enrichment analysis (GSEA) with curated pathway annotations to visualize signaling signatures derived from the proteomic content of circulating telomere Rivers (Figure 3a), as derived from KEGG and Gene Ontology Biological Processes. Pathway-level connectivity to stem-related WNT, Notch, and Runx1 signaling is shown. Arrows represent directionally inferred activation, based on curated datasets (STRING confidence score > 0.7).

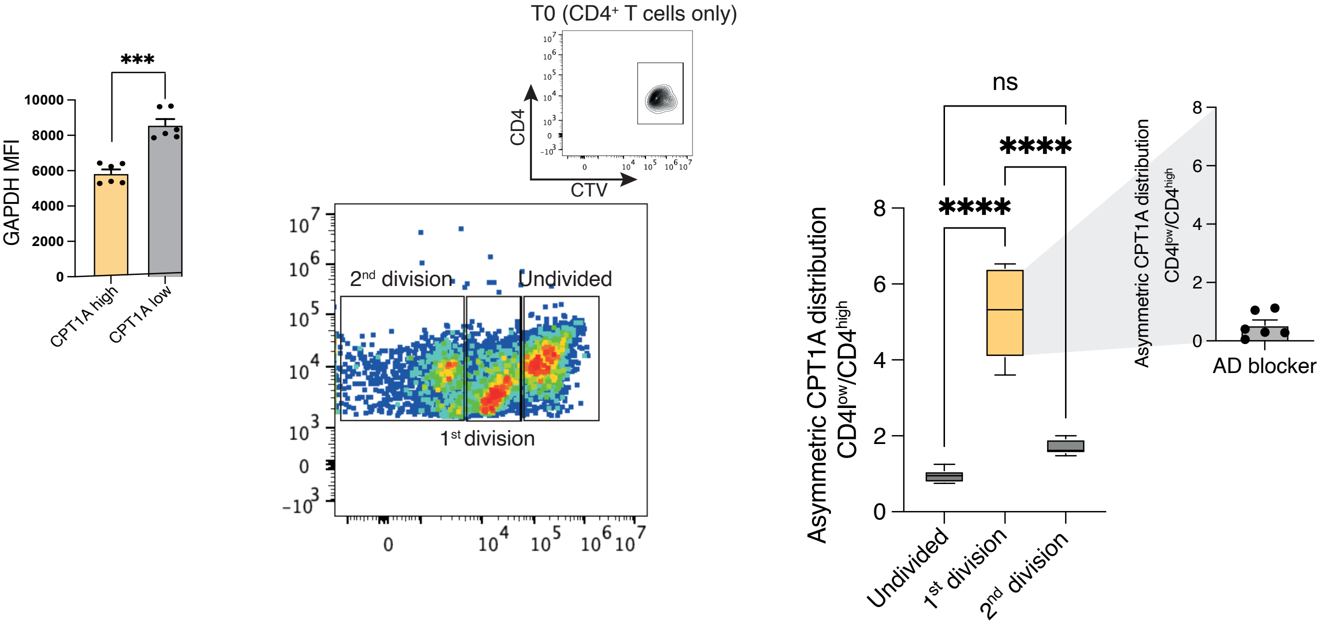


**Supplementary information 5:** Asymmetric capacity is acquired during the first round and lost in subsequent rounds of T cell division. Representative CTV FACS plot as shown in Figure 3d (left) and quantification of CPT1A (right) asymmetric distribution in $CD4^{low}$ and in $CD4^{high}$ T cells, expressed as fold change across different rounds of T cell division. T0, CTV staining before cell division. GAPDH levels in Cpt1a high vs low stem like T cells is shown (left). An additional graph showing the lack of asymmetric CPT1A distribution in the presence of the anti-LFA1 blocking antibody is also included (top right).

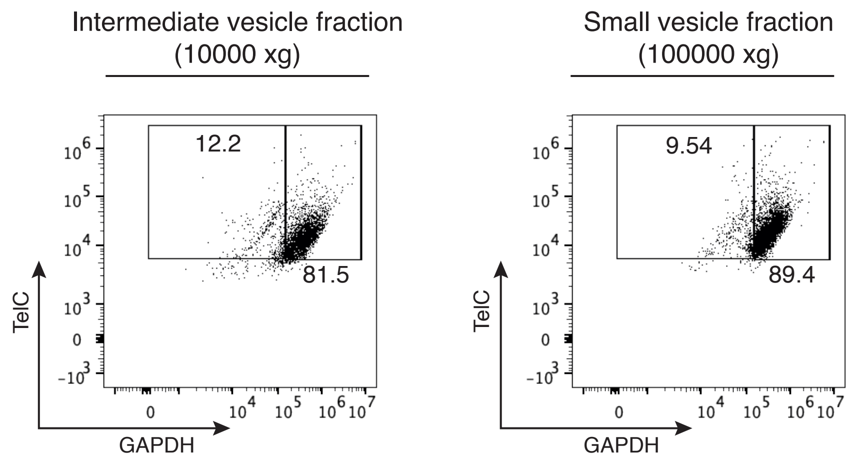


**Supplementary information 6:** Further vesicle fractions obtained by sequential centrifugations (10000 xg, 100000 xg). The larger River fraction (3000 xg) is shown in Figure 3f.

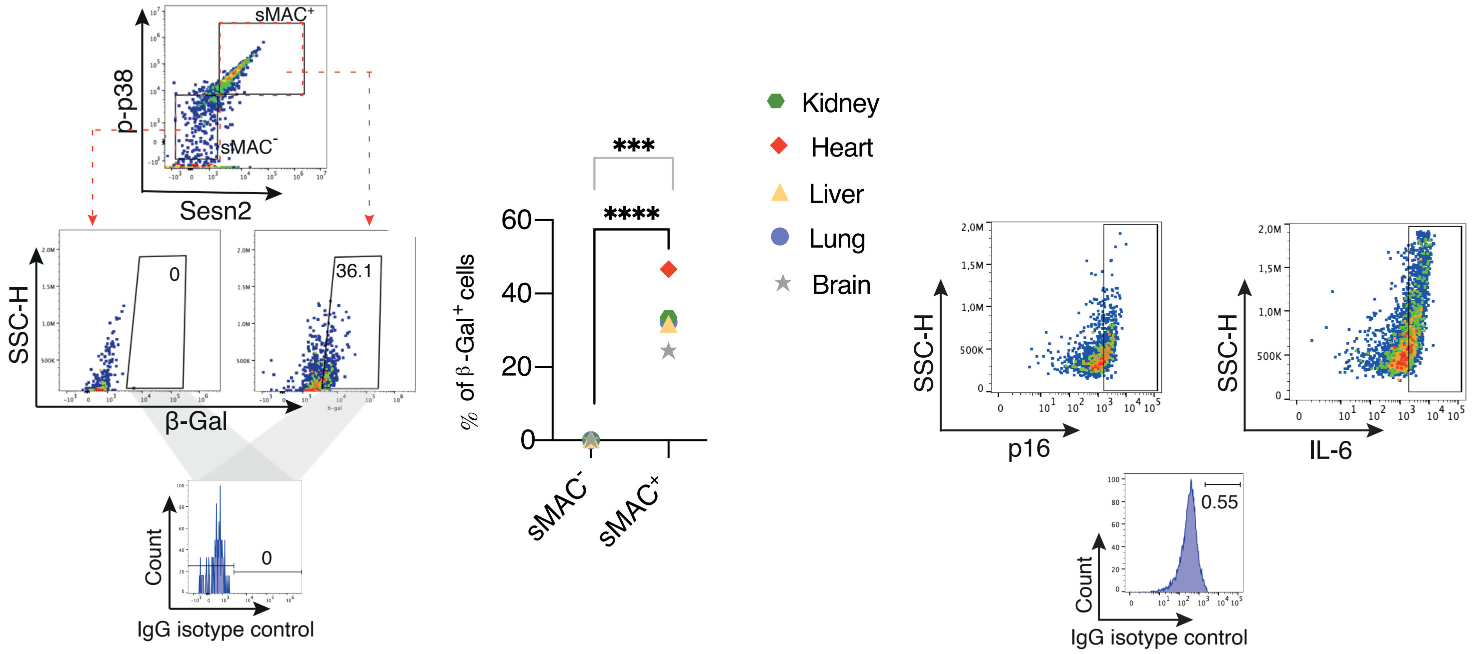


**Supplementary information 7:** Representative FACS plots (brain) and quantification of the senescence marker β-Galactosidase in sMAC+ and in sMAC- cells of different mouse organs (left).
Note that β-Galactosidase is only found among sMAC+ cells in the different organs.
Representative FACs plots for p16 and IL-6 (brain) referred to Extended Data Fig. 10 are shown (right).
IgG isotype controls are included for comparison.
